# A gelation transition enables the self-organization of bipolar metaphase spindles

**DOI:** 10.1101/2021.01.15.426844

**Authors:** Benjamin A. Dalton, David Oriola, Franziska Decker, Frank Jülicher, Jan Brugués

**Affiliations:** Max Planck Institute of Molecular Cell Biology and Genetics, 01307, Dresden, Germany; Center for Systems Biology Dresden, 01307 Dresden, Germany; Max Planck Institute for the Physics of Complex Systems, 01187, Dresden, Germany; Cluster of Excellence Physics of Life, TU Dresden, 01307 Dresden, Germany

**Author notes:** Department of Physics, Freie Universität Berlin, 14195, Berlin, Germany (Benjamin A. Dalton). EMBL Barcelona, 08003, Barcelona, Spain. ETH Zürich, D-BSSE, 4058, Basel, Switzerland. These authors contributed equally (Benjamin A. Dalton, David Oriola, Franziska Decker).

## Abstract

The mitotic spindle is a highly dynamic bipolar structure that emerges from the self-organization of microtubules, molecular motors, and other proteins. Sustained motor-driven poleward flows of short dynamic microtubules play a key role in the bipolar organization of spindles. However, it is not understood how the local activity of motor proteins generates these large-scale coherent poleward flows. Here, we combine experiments and simulations to show that a gelation transition enables long-ranged microtubule transport causing spindles to self-organize into two oppositely polarized microtubule gels. Laser ablation experiments reveal that local active stresses generated at the spindle midplane propagate through the structure thereby driving global coherent microtubule flows. Simulations show that microtubule gels undergoing rapid turnover can exhibit long stress relaxation times, in agreement with the long-ranged flows observed in experiments. Finally, we show that either disrupting such flows or decreasing the network connectivity can lead to a microtubule polarity reversal in spindles both in the simulations and in the experiments. Thus, we uncover an unexpected connection between spindle rheology and architecture in spindle self-organization.

## Introduction

Cells use active cytoskeletal structures to perform essential processes such as cell division, cell motility, force generation, and morphogenesis (*Mitchison and Cramer, 1996, Howard, 2001, Heisenberg and Bellaíche, 2013, Cross and McAinsh, 2014*). Cytoskeletal structures are made up of dynamic filaments that are cross-linked by molecular motors and rely on constant energy consumption for their assembly and function. Molecular motors drive the organization of cytoskeletal structures by actively clustering, pulling, and sliding filaments apart (*Miyamoto et al., 2004,Tolić-Nørrelykke, 2008, Dumont and Mitchison, 2009, Howard, 2009, Sanchez et al., 2012*). This activity often leads to the emergence of large-scale flows within the structures that can transport material (*Mayer et al., 2010*), exert force (*Boukellal et al., 2004*), or establish a gradient of filament polarity (*Brugués et al., 2012*). Although the emergence of flows in active matter has been successfully captured by hydrodynamic theories (*Joanny et al., 2007, Marchetti et al., 2013, Prost et al., 2015*), the microscopic origins of flows and their relation to the rheology and architecture of cellular structures remains poorly understood.

A prominent example of active large-scale flows is the poleward movement of microtubules observed in the metaphase spindle (*Mitchison, 1989, Maddox et al., 2002, Ganem and Compton, 2006*). Spindles are self-organized molecular machines responsible for the segregation of sister chromatids during cell division. They are made of dynamic microtubules that are continuously transported towards the poles by molecular motors in a process known as poleward flux (*Mitchison, 1989, Maddox et al., 2002*). In *Xenopus laevis* egg extract spindles, this flux depends on the activity of Eg5, a kinesin motor that can slides antiparallel microtubules (*Miyamoto et al., 2004, Kapitein et al., 2005, Uteng et al., 2008, Striebel et al., 2020*). As a consequence, microtubule flows should depend on the spatial distribution of antiparallel microtubule overlaps in the structure. Laser ablation measurements revealed that the density of antiparallel microtubule overlaps forms a pronounced gradient in the spindle, with maximum antiparallel overlap in the spindle mid-plane (*Brugués et al., 2012*). However, flows are remarkably constant throughout the structure (*Yang et al., 2008*), which raises the question of how a gradient of antiparallel microtubule overlaps can result in a flow that is spatially homogeneous.

Recent work *in vitro* demonstrated that mixtures of stabilized microtubules and motors can generate active flows that are independent of the spatial distribution of antiparallel microtubule overlaps (*Fürthauer et al., 2019*). It was suggested that these flows can arise from the high degree of connectivity introduced by molecular motors. However, in contrast to stabilized mixtures, microtubules in the spindle are constantly nucleated throughout the structure (*Oh et al., 2016, Decker et al., 2018, Rieckhoff et al., 2020*) and turnover rapidly (*Brugués et al., 2012, Redemann et al., 2017, Rieckhoff et al., 2020*). In particular, microtubule nucleation is an auto-catalytic process (*Ishihara et al., 2014, Oh et al., 2016, Decker et al., 2018*), which leads to microtubule structures that are genuinely different from the ones obtained in other simplified *in vitro* systems (*Sanchez et al., 2012, Roostalu et al., 2018, Fürthauer et al., 2019*). The main two reasons being that microtubule nucleation in the spindle requires the presence of pre-existing microtubules (*Petry et al., 2013, Ishihara et al., 2014, Oh et al., 2016, Decker et al., 2018, Kaye et al., 2018, David et al., 2019, Verma and Maresca, 2019*) and that newly nucleated microtubules are aligned with respect to their mother microtubules (*Decker et al., 2018, Thawani et al., 2019*). In the absence of microtubule transport, these waves of autocatalytic microtubule nucleation and growth lead to the formation of monopolar spindles (*Decker et al., 2018*), with microtubule plus-ends predominantly point outwards, away from the center of the structure, opposite to what is found in spindles. How the interplay between microtubule active flows and autocatalytic microtubule waves establish the correct spindle architecture remains unknown.

Here we combine experiments and simulations to show that a gelation transition enables long-ranged active flows in *Xenopus* egg extract spindles. This transition leads to the self-organization of two polar, interpenetrating, and mechanically distinct microtubule gels of opposite polarity. We show that motor sliding activity is localized in the spindle midplane region, where the two gels maximize antiparallel overlaps, and that motor activity continuously pushes the two gels apart. Simulations of highly cross-linked active microtubule networks reveal that despite continuous microtubule turnover, microtubule gels can sustain long-ranged steady-state flows and exhibit an elastic-like response at time scales greater than the microtubule lifetime. Finally, we show that disrupting microtubule flows or the network connectivity leads to the reversal of microtubule polarity in spindles, illustrating a functional relevance of gelation in the proper organization of metaphase spindles, and elucidating an unexpected connection between gelation, microtubule nucleation, and microtubule transport.

## Results

### Local sliding of antiparallel microtubules cannot account for the poleward flux

The local microtubule-sorting hypothesis implies that the poleward motion of microtubules throughout the spindle should depend spatially on the antiparallel microtubule overlap density (*Walczak et al., 1998, Uteng et al., 2008, Brugués et al., 2012*). To test this hypothesis, we quantified microtubule transport and the density of antiparallel overlaps throughout the spindle structure. To focus on microtubule transport alone we inhibited dynein activity, which is known to slightly affect poleward flux at the poles and to control spindle shape (*Uteng et al., 2008, Yang et al., 2008, Oriola et al., 2020*) (see Fig. 1, Fig. S1A and Movie S1). To this end, we treated spindles with p150-CC1 (*Mayer et al., 1999*) which is known to disrupt pole formation, causing microtubule alignment along the spindle long axis (see Fig. 1A) (*Gaetz and Kapoor, 2004*). Dynein inhibition caused only a minor effect on spindle architecture (see Fig. S1B-D). To visualize the motion of individual microtubules in spindles, we used singlemolecule speckle microscopy (*Yang et al., 2008, Brugués et al., 2012*) (Fig. 1B, C and Movie S2). Speckle trajectories showed two distinct populations (Fig. 1D) moving coherently towards the spindle poles. To compare the profile of microtubule velocities to the antiparallel microtubule overlaps, we quantified the density distribution of the two microtubule populations (Fig. 1E), revealing two interpenetrating networks, in agreement with previous studies (*Yang et al., 2008*). Computing the density of microtubule overlaps using the two distributions, we found that the number density of antiparallel overlaps was maximal at the spindle center and decreased by ~ 90 % at the poles (see Fig. 1F). In contrast, microtubule transport was found to be constant throughout the spindle with an average value of 2.5 ± 0.4 μm/min (n = 549 speckles from 10 spindles, mean ± SD, Fig. 1F and S1A) in agreement with Ref. (*Yang et al., 2008*). To ensure that speckle microscopy provides an accurate readout of microtubule polarity, we compared the relative difference between the two microtubule populations (Fig. 1G, circles) to the microtubule polarity profile obtained by laser ablation (Fig. 1G, squares) (see Material and Methods and Ref. (*Brugués et al., 2012*)). The two approaches were in agreement, confirming that the direction of speckle transport reveals microtubule polarity. We thus conclude that microtubule transport velocity is independent of local antiparallel overlap density (i.e. microtubule polarity see Fig. S2) and thus cannot be explained by the local sorting of microtubules alone.

**Figure 1:**
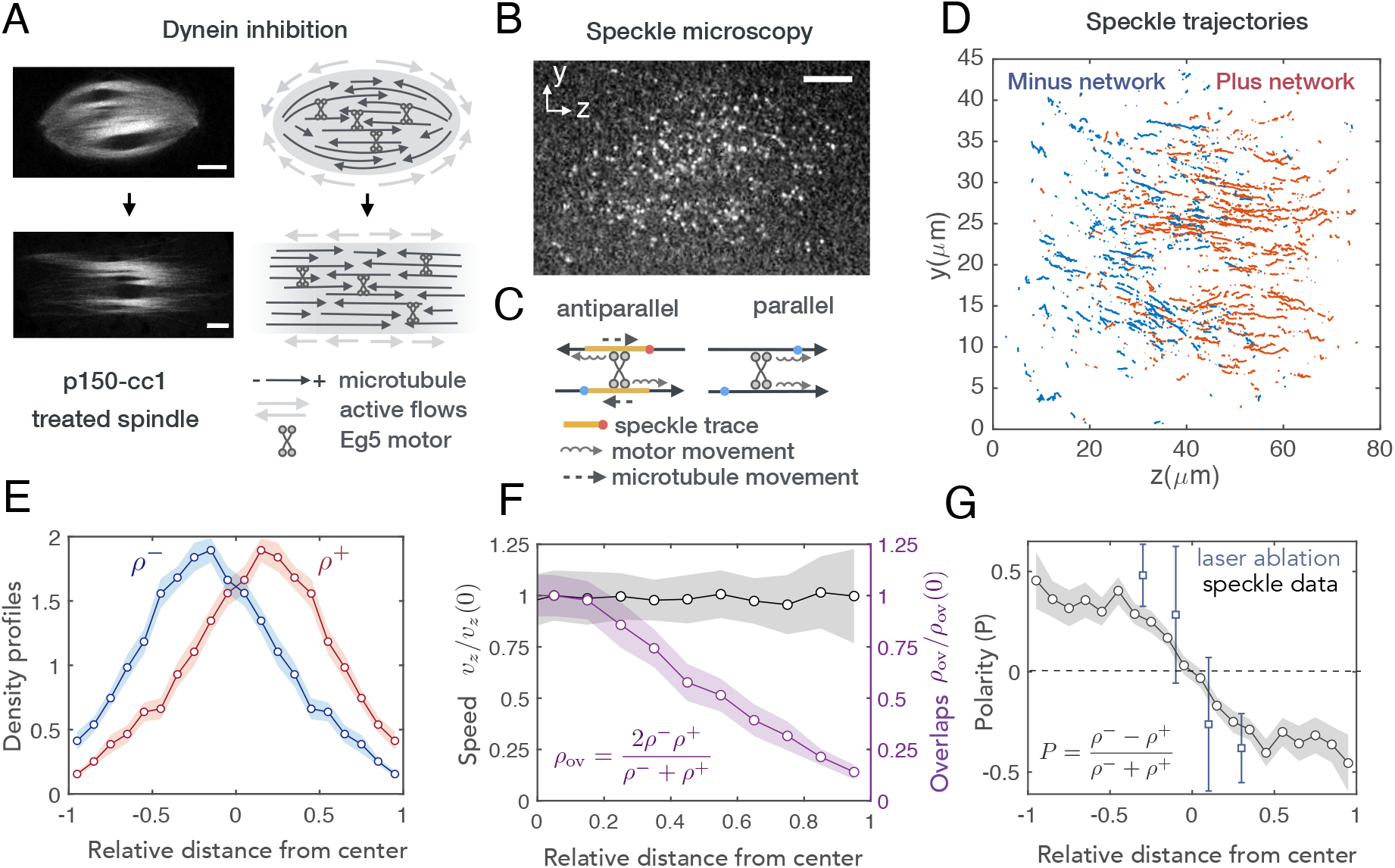
Local sorting of antiparallel microtubule overlaps is not sufficient to explain poleward microtubule transport. (A) Fluorescent spindles labeled with atto565 pig tubulin and corresponding schematics of spindle architecture for the control and dynein inhibited cases (Scale bar 10 *μ*m). (B) Speckle microscopy (~ 1 nM atto565 frog tubulin) is used to measure microtubule transport (see Movie S2). Scale bar: 10 *μ*m. (C) Speckles move in antiparallel overlaps due to the action of Eg5 motors but they remain static in parallel overlaps. (D) Speckle trajectories are tracked and classified according to their direction of motion. Speckles whose longitudinal velocity was smaller than 0.2 *μ*m/min were discarded from the analysis (~ 4 %). Total timelapse is 210 s. (E) Number density of the two microtubule populations (mean ± SD, *n* = 10 spindles). (F) Normalized value of the averaged velocity profiles of the plus and minus networks compared to the normalized number density of antiparallel overlaps (see Appendix, Experimental Methods). Eg5 can only generate forces in the spindle center where antiparallel overlaps are enriched. (mean ± SD, *n* = 10 spindles). (G) Comparison between polarity profile obtained from the speckle data (mean ± SD, *n* = 10 spindles) and the polarity profile obtained using laser ablation (*n* = 25 cuts, mean ± SD). This result provides direct evidence that the direction of movement of the speckles is a readout of microtubule polarity.

### Microtubule transport is driven by long-ranged stress propagation throughout the spindle

Our speckle analysis shows that local sorting is not sufficient to explain microtubule transport in spindles. One possible explanation for the polarity-independent flows observed throughout the spindle is that the local stresses generated by Eg5 in the region of antiparallel overlaps, which concentrate around the spindle midplane, are propagated over long distances. To test this hypothesis, we designed an assay combining fluorescent speckle microscopy and laser ablation (see Fig. 2A and Material and Methods). We reasoned that if Eg5 activity is mainly restricted to the spindle midplane and propagates throughout the structure, disconnecting a region of the spindle that is far from the midplane using laser ablation should considerably reduce microtubule transport—as measured by the motion of speckles— within the disconnected region. (Fig. 2A, a). Alternatively, if Eg5 is acting homogeneously in the structure via local sorting, microtubule motion in the disconnected region should remain unaffected (Fig. 2A, b). A key aspect of this perturbation is that the poleward-moving speckles that survive after the laser cut would correspond to intact antiparallel overlaps. This is because cut microtubules only depolymerize from the newly created plus end, whereas the new minus ends remain stable (see Fig. 2B).

**Figure 2:**
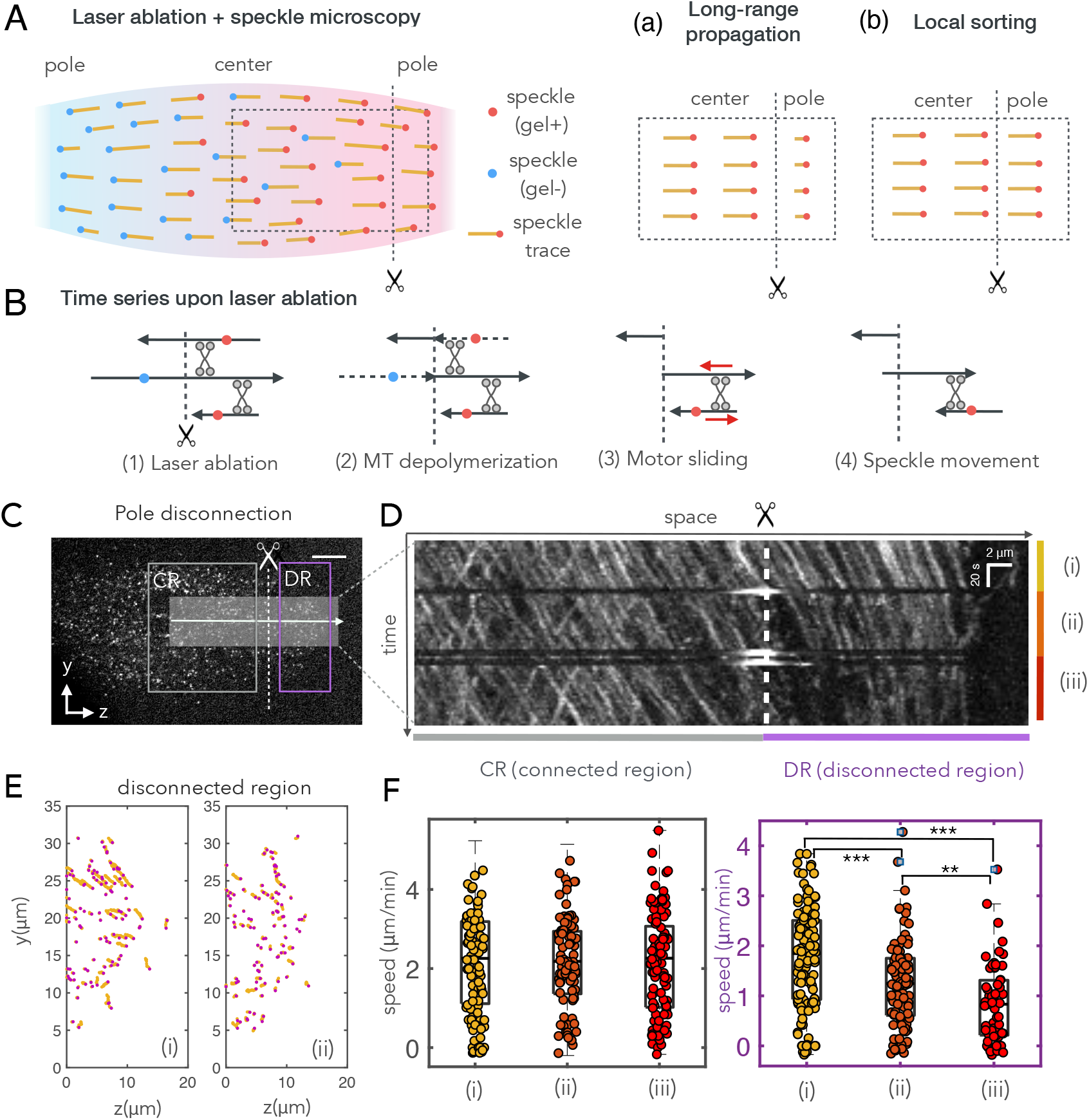
Microtubule transport is a consequence of long-ranged stress propagation mediated by Eg5. (A) A combination of laser ablation and speckle microscopy techniques are used to distinguish between two scenarios: (a) Eg5 generates stresses at the center of the spindle and they propagate across the structure. Speckles stop in the disconnected region after laser ablation whereas speckles in the rest of the spindle remain unaffected. (b) Eg5 generates local stresses throughout the structure and speckles are not affected in the disconnected region after the cut. (B) Laser ablation does not affect microtubule transport: Laser ablation induces the depolymerization of some microtubules and the consequent unbinding of some motors. However, the speckles that survive after laser ablation in the disconnected pole (right side in the diagram) can still move if antiparallel overlaps are present in the disconnected region. Hence, the laser ablation method does not affect the movement of the remaining speckles in the disconnected region. (C) Fluorescent speckle image before laser ablation. Scale bar: 10 *μ*m. CR: connected region, DR: disconnected region. The arrow denotes the direction along which the kymograph in (D) is studied and the width of the gray region denotes the averaged region. (D) Kymograph from (C) showing how speckle trajectories stop in the disconnected region after consecutive laser ablation events. This result validates the scenario (a). (E) Speckle trajectories before (time interval (i)) and after (time interval (ii)) the first laser ablation during 40 s (see Movies S4 and S5). (F) On-axis velocity of speckles during the time intervals (i), (ii), and (iii) in the connected (CR) and disconnected (DR) regions. A progressive reduction in speckle velocity is observed in the disconnected region while the speckle velocity remains unaffected in the connected region. Only speckles with > −0.2 *μ*m/min were considered in the analysis. A reduced subset of 100 data points is shown in the disconnected region case for the sake of clarity. ** *p* < 0.01 and *** *p* < 0.001. Squares indicate outliers. The test used was a two-sample t-test.

We achieved complete disconnection of a spindle pole using laser ablation by producing consecutive cuts across the body of the spindle (Fig 2A-C and Movie S3). The microtubule velocity was significantly reduced after ablation in the disconnected region (DR) from 1.8 ± 0.1 μm/min (T1, *n* = 122 trajectories, median ± SEM) before the cut to 0.8 ± 0.1 μm/min after a series of cuts (T3, *n* = 61 trajectories, mean ± SEM) (see Fig. 2D,E,F and Movies S4-S6). In contrast, microtubule velocity remained unaffected in the connected region (CR) (see Figs. 2D, E, F and Movies S7-S9). We obtained similar results in other replicas of the experiment (see Fig. S3 and Movies S10 and S11). The reduction in transport in the disconnected region after ablation indicates that microtubule flux in that region is not a consequence of local microtubule sorting or treadmilling at spindles poles as previously proposed (*Rogers et al., 2005, Needleman et al., 2010, Oriola et al., 2018*). Instead, local stresses generated by Eg5 propagate throughout the structure from the spindle midplane region to the poles, driving microtubules poleward.

### The spindle is composed of two interacting and interpenetrating microtubule networks of opposite polarity

We wondered how the two microtubule populations in the spindle can sustain long-range stress propagation, especially given the short length of microtubules compared to the spindle size and their fast turnover (*Needleman et al., 2010, Brugués et al., 2012*). To gain insight into the material properties of the two networks, we measured the fluctuations of speckle pairs in dynein inhibited spindles (Fig. 3A-D) and performed two-point correlation analysis on these fluctuations (Fig. 3E-G) (see Material and Methods), (*Crocker et al., 2000, Levine and Lubensky, 2001, Lau and Lubensky, 2009, Brugués and Needleman, 2014*). We calculated the two-point correlations along the transversal (y) and longitudinal (z) axis of the spindle, as a function of the distance between speckles *r*. We classified the speckles according to their direction of movement and considered three different cases: pairs belonging to the same network, pairs from different networks, and pairs taken indiscriminately from either population (Fig. 3E, F, and G, with yellow, gray and blue colors, respectively). Transversal correlations decayed as ~ 1/*r* (Fig. 3E) in agreement with correlations expected for a continuum material (*Crocker et al., 2000, Levine and Lubensky, 2001, Brugués and Needleman, 2014*). Strikingly, when we considered either speckle pairs from a single network or speckle pairs from different networks, we did not observe any decay in the longitudinal correlations (Fig. 3F, yellow and gray). These results suggest that the two networks behave as rigid-like materials along the spindle long-axis while being deformable along the short-axis. The fluctuations of speckle pairs coming from the same network were positively correlated, implying a coherent stress transmission across each network. In contrast, the fluctuations of speckle pairs from different networks were negatively correlated, suggesting that Eg5 acts on the two networks pushing them apart. When we combined all the speckles from both networks, the corresponding correlations decayed with distance (Fig. 3F, blue). However, this dependence is a consequence of a microtubule polarity gradient in the spindle, that dictates the probability that two random speckles belong to the same or opposite networks. Finally, we reasoned that if the two networks fluctuate as whole entities, shear fluctuations would not be present. We quantified correlations in shear fluctuations by calculating the propagation of longitudinal fluctuations as a function of the transversal direction (Fig. 3G). Remarkably, correlations did not decay in space, further confirming that each network fluctuates as a whole. Together, these results suggest that the spindle is composed of two interpenetrating rigid-like microtubule networks of opposite polarity that are coupled at the central overlap region and driven apart by Eg5 activity.

**Figure 3:**
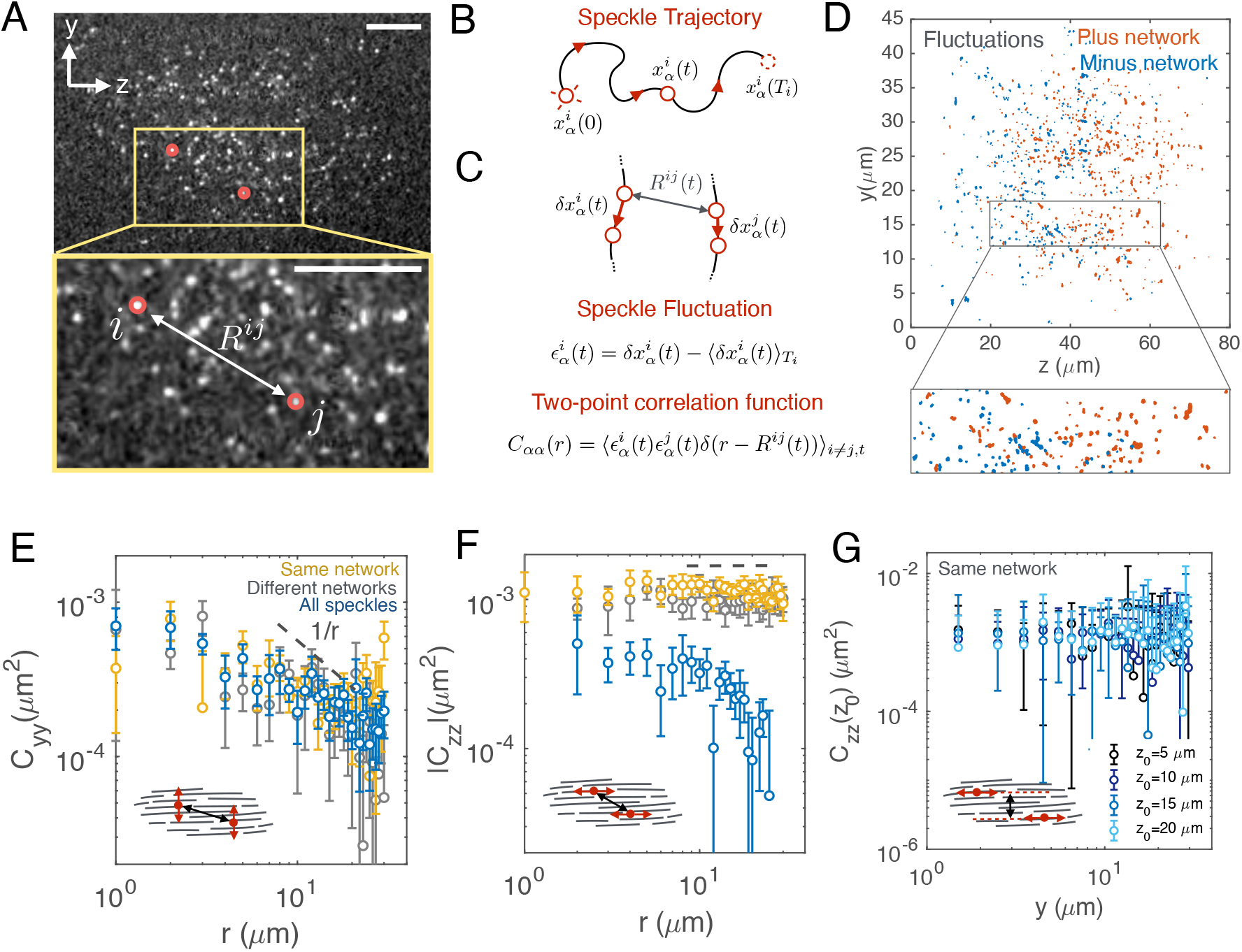
Two-point correlations reveal two interacting rigid-like microtubule networks in the spindle. (A) Fluorescently labelled tubulin speckles in a control spindle. Zoomed region: Speckle pair (*i,j*) sep arated by a distance Rij. Scale bar: 5 *μ*m. (B) Speckle trajectory tracked from its appearance at *t* = 0 and disappearance at time *t* = *T_i_*. (C) Speckles pair trajectories and positions at two different times separated by Δ*t*. (D) Speckle fluctuations and classification according to their mean velocity along the spindle long-axis *z*. Speckles moving towards the right are coloured in orange, while speckles moving towards the left are coloured in blue. Total timelapse is 210 s. (E), (F) and (G) Log-log plots of the two-point correlations for dynein-inhibited spindles (*n* = 9), *C_yy_* (r) and *C_zz_* (*r*) of speckles classified in the same network (yellow), different networks (gray) or not classified (blue) (see Appendix: Experimental Methods). Transversal fluctuations decay and are decoupled between the two networks, while longitudinal correlations are coupled between networks and do not decay within networks. (G) Shear correlations *C_zz_* (*y*; *z*_0_) for *z*_0_ = 5,10,15, 20 *μ*m (from dark to light blue) (see Appendix: Experimental Methods).

### A physical model shows that a gelation transition enables long-range propagation of motor-generated stresses

Mixtures of filaments and motors can behave as fluids or gels depending on the filament connectivity. Above a critical connectivity threshold, the mixture undergoes a percolation transition where all components are interconnected within a cluster that spans the entire system (*Broedersz and MacKintosh, 2014, Alvarado et al., 2017*). Within this regime, the system can behave as a gel even if its components continuously exchange. This is true provided that the connectivity is maintained above the rigidity threshold (*Thorpe and Duxbury, 1999, Ehrlicher et al., 2015, Alvarado et al., 2017*). Despite of the fact that gelation has been intensely studied in actin systems (*Stossel et al., 1987, Field et al., 2011, Alvarado et al., 2013, Lee and Pruessner, 2016, Alvarado et al., 2017*) it has remained largely unexplored in microtubule systems. To test whether a gelation transition underlies the long-range propagation of stresses in the spindle we turned to large-scale simulations (see Appendix: Simulation Methods for details). To simplify the two overlapping microtubule networks observed in the spindle, we simulated the dynamics of one filament network and substituted the other for a pinning field structure of fixed, polar aligned filaments. The length of the fixed arrangement of filaments controlled the region of antiparallel overlaps representing the spindle midplane (Fig. 4A,B, red region). Simulations included filament nucleation and turnover, with the orientation of new filaments being antiparallel to the pinning field, as well as active force generation by transient motors that walked along the filaments mimicking Eg5 (see Fig. S4 for a schematic overview). Eg5 performed both the role of a crosslinker in parallel overlaps and the role of a motor in antiparallel overlaps. To ensure a high filament density and reproduce the conditions for microtubule orientation in the spindle, we confined filaments to narrow channels with strongly repulsive walls that forced the filaments into a dense polar phase (see Fig. S5). This minimal system enabled us to study how localized active stresses can propagate through cross-linked filament networks.

**Figure 4:**
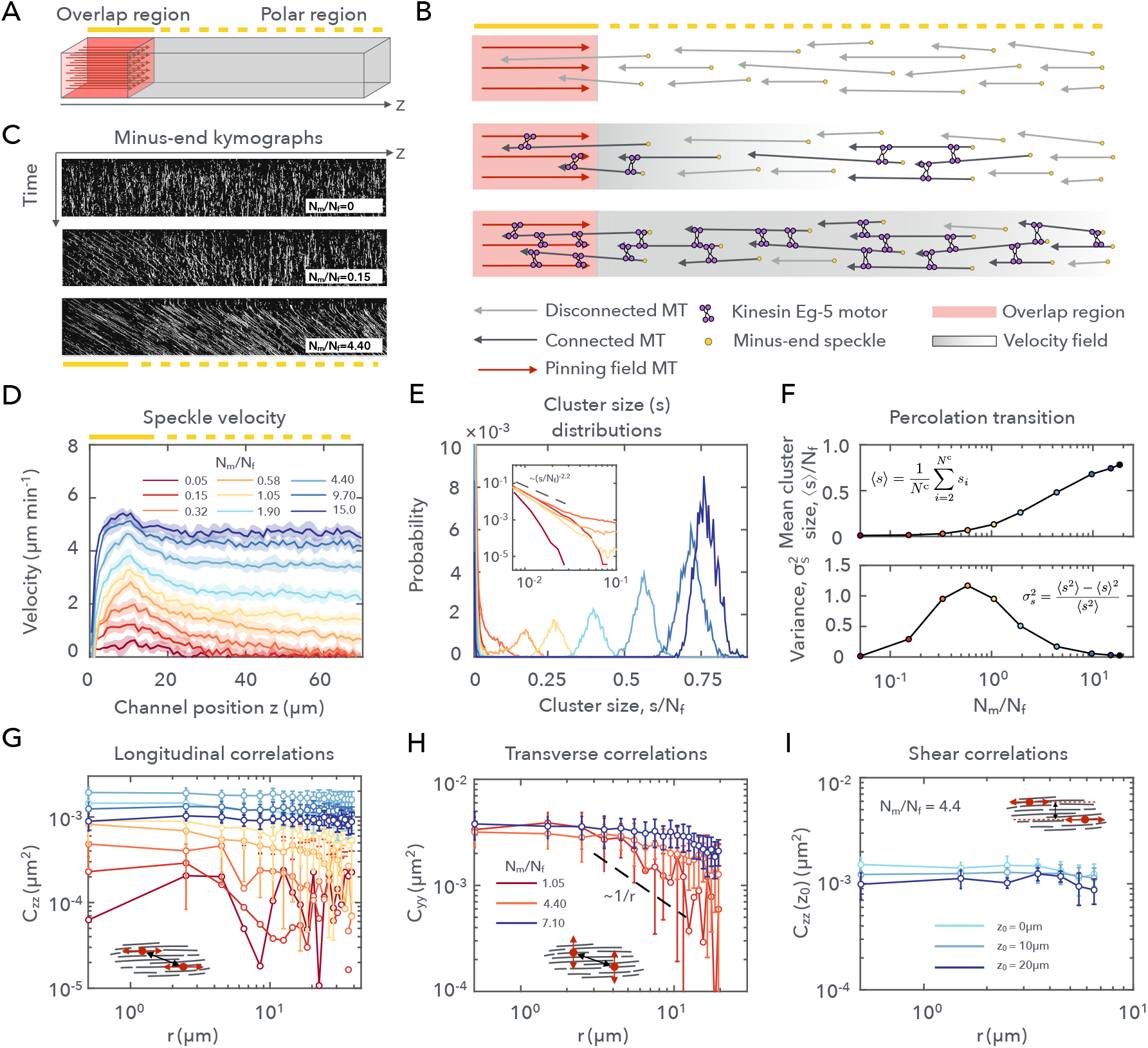
A gelation transition enables long-ranged flows in filament networks. Computer simulations of filament networks in a pinning-field channel. (A) The pinning field substitutes for one of two filament networks. Schematic shows the pinning-field channel simulation environment. (B) Conceptual overview for simulations: dynamic microtubules nucleate throughout the channel, restricting antiparallel overlap to the pinning field region. Increasing Eg5 concentration increases the filament connectivity, enabling long-ranged flow. Solid walls constrain filament orientations. (C) Kymograph for minus-end speckles showing a line section along the channel long-axis for three different Eg5 concentrations (Movie S12). (D) The length-scale of the filament velocity profiles increases with increasing Eg5 concentration (figure legend shows color scheme for (D), (E), and(G)). (E) Distributions *P*(*s*/*N_f_*) of filament cluster size *s/N_f_* indicate a percolation transition in filament networks. Distributions are calculated as a function of Eg5 concentration. (inset) Near the percolation threshold, the distribution approaches a power-law distribution with exponent ~ 2.2, in agreement with percolation theory. (F) Mean and the normalized variance for the cluster size distributions in (E). (G) Longitudinal two-point fluctuation correlations reveal the onset of rigid-like behavior in gelated networks. Correlations are shown for simulations in (D), (E) and (F). (H) Transverse two-point correlations decay for high Eg5 concentrations. A wider simulation domain is used to probe transverse fluctuations (see Material and Methods: simulating active, long-range flows). (I) Shear correlations for gelated networks are also constant over a range of *z_0_* values. Channel geometry is the same as in (H).

To study the effect of the degree of filament connectivity on filament transport, we systematically varied the number of motors in the system (Fig. 4C and Movie S12). We found that for motor concentrations below 0.5 motors/filament, flows decayed abruptly beyond the overlap region with a length scale close to the mean length of a filament (Fig. 4D). However, above this concentration, we observed the emergence of coherent flows that spanned the entire channel (up to a range ~ 10 times larger than the mean filament length). To test if the emergence of these long-range flows corresponds to the onset of a gelation transition, we measured the connectivity within the cross-linked filament networks and calculated the distribution of cluster sizes as a function of motor concentration (Fig.4E). At approximately 0.5 motors/filament we observed the emergence of a distinct peak in the cluster size distribution accompanied by a maximum in the cluster size variance (Fig. 4F), indicating that the system undergoes a percolation transition. The peak in the cluster size distribution indicates the formation of a filament cluster that spans the entire length of the channel, enabling the long-range propagation of stresses generated at the pinning field. The size of the dominant cluster increases as a function of motor concentration and saturates for highly-crosslinked networks (10-15 motors/filament, similar to estimated values for spindles in Refs. (*Brugués and Needleman, 2014, Furthauer et al., 2019*)).

We next tested whether the gelation transition accounted for the material properties of the spindle by performing two-point microrheology measurements in our simulations. In the gelated conditions, these measurements showed flat long-ranged correlations along the channel and a ~ 1/r decay of the correlations transverse to the channel (see Fig. 4G, H), in agreement with experiments (Fig. 2E, F, yellow curves). Additionally, the flows along the channel exhibit no shear fluctuations (Fig. 4I) consistently with the experimental observations (Fig. 2G). Altogether, our simulations show that a gelated polar network is sufficient to sustain the long-range flows observed in spindles.

### Polar gels behave as viscoelastic fluids with a relaxation time that can be larger than the filament turnover time

Our experiments and simulations suggest that the gelation of microtubule networks in spindles results in the emergence of two interpenetrating gels. However, the binding and unbinding of motor proteins and the rapid microtubule turnover should fluidize the gels at long timescales (*Shimamoto et al., 2011*). To understand the rheological properties of polar filament gels, we oscillated the pinning field in our simulations and studied the response of a filament probe trapped in a harmonic potential far from the pinning field (see Fig. 5A-C, see SI). For simplicity, motor activity was turned off for the microrheology measurements to avoid active contributions to the rheological properties of the network. We measured the response of the probe for oscillation frequencies in the range ~ 0.003 — 0.1 s^−1^. This method mimics previous microrheological measurements using a combination of rigid and flexible microneedles in spindles (*Shimamoto et al., 2011*), although in these measurements it was not possible to distinguish between the two gels. To characterize the rheological properties of the network we calculated the in-phase and out-of-phase response of the network deformation, *Z′* and *Z″* respectively, to the periodic forcing (see appendix: AMR and calculating sinusoidal response and Fig. S6A). We found that in the moderate crosslinked regime, and at high frequencies (0.1 s^−1^), the response of the trapped filament is in-phase to the network deformations, implying that the gel behaves as an elastic solid (Fig. 5C, left). In contrast, for low frequencies (0.008 s^−1^) the response is out-of-phase and the system behaves as a viscous fluid (Fig. 5C, right). We observed that the crossover frequency from elastic to fluid response varied with the crosslinker concentration (Fig. S6A). However, by rescaling *Z′, Z″*, and the frequency with respect to the corresponding values at the crossover point for different crosslinker concentrations, we found that all the curves collapse following a simple Maxwell model with a single relaxation timescale (see Fig. 5E). The relaxation time increases with increasing crosslinking and saturates for ~ 10 crosslinks/MT (Fig. 5E, inset). Strikingly, for high crosslinker concentrations, the relaxation time is ~ 80 s, which is approximately 3 to 4 times larger than the mean filament lifetime. This relaxation time scale depends on the microtubule length distribution and on the crosslinker binding/unbinding kinetics, such that both increasing the mean filament length and decreasing the motor unbinding rate leads to an increased time scale, and vice versa (Fig. S6B). In summary, filament gelation can lead to elastic-like responses over minute time scales despite the rapid filament and crosslink turnover.

**Figure 5:**
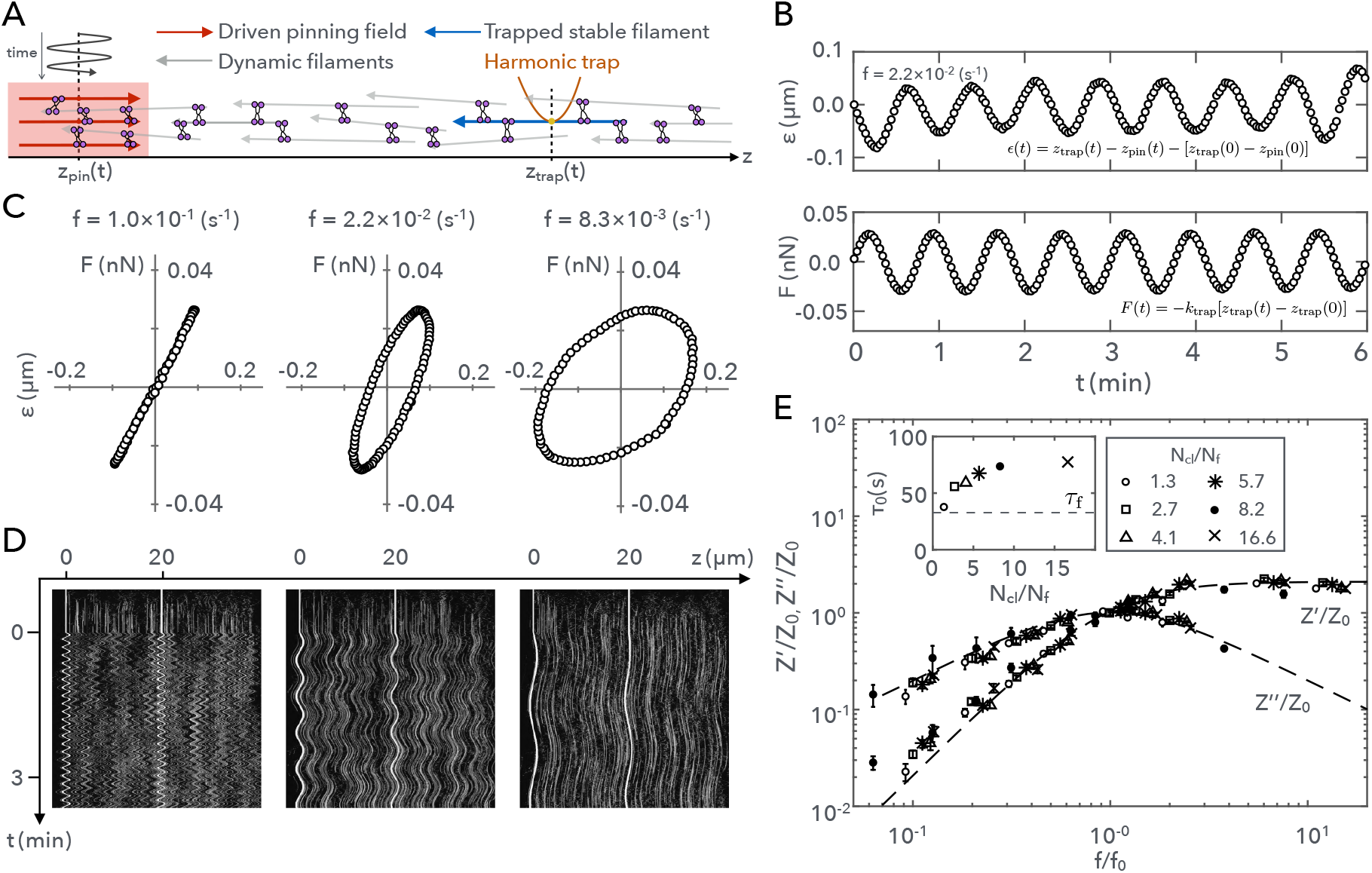
The viscoelastic relaxation of the gel is larger than the filament lifetime. Active microrheology (AMR) measurements for viscoelastic response in simulated filament networks. (A) Schematic showing AMR setup, including oscillating pinning field and trapped probe filament. The permanent probe filament is confined to a harmonic trap in *x, y* and *z*, set at *z*_trap_ = 20 *μ*m. The pinning field oscillates according to *z*_pin_(*t*) = *z*_0_ sin(2*πft*), with *z*_0_ = 0.75 *μ*m. Network connectivity, due to passive crosslinking, enables stress propagation from the pinning field to the trapped probe filament. (B) Network deformations are driven by pinning field oscillations and resistance of the trapped filament. Network deformations given by *ϵ*(*t*) = *z*_trap_(*t*) – *z*_pin_(*t*) – *L*_0_, where *L*_0_ = 20*μ*m. Trajectories of probe filaments indicate the force response. (C) Phase plots show that contributions from viscous and elastic responses vary depending on driving frequency. Phase response is shown for three frequencies (*N*_cl_/*N*_f_ ≈ 4). (D) Kymographs for gelated networks show the probe moving in response to the driven pinning field. Driving frequencies are the same as in (C). (E) Viscoelastic transition times vary depending on crosslinker concentration. The plot shows rescaled material viscoelastic response over a range of crosslinker concentrations. Fit for a Maxwell model included for *Z′/Z*_0_ = *A′* (*f/f*_0_)^2^/(1 + (*f*/*f*_0_)^2^) and *Z′*/*Z*_0_ = *A″*(*f/f*_0_)/(1 + (*f*/*f*_0_)^2^), where *A′* = 2.10 and *A″* = 2.01. (inset) Viscoelastic transition times increase with increasing crosslinker concentration, saturating at ~ 3-4 τ_MT_.

Our rheological measurements allowed us to estimate the effective viscosity and elastic modulus of the filament gel as a function of the crosslinker concentration (see Fig. S6C, D, and Appendix: Calculating network viscosity and hydrodynamic length). At long time scales, the range at which flows propagate is given by the hydrodynamic length scale 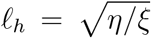, where *η* is the effective viscosity of the gel and ξ is the friction with the solvent (see Appendix). For moderately crosslinked networks (~ 5 crosslinkers/MT) we obtain a hydrodynamic length scale *ℓ_h_* ~ 300 μm which is much larger than the channel length. Thus, we conclude that long-range stress propagation in the spindle is a consequence of a large relaxation time, despite fast microtubule turnover, which emerges from the gelation transition.

### A gelation transition enables the correct bipolar spindle architecture

Our results show that above the gelation transition point, flows become long-ranged and independent of the filament polarity profile, whereas below the transition point flows decay over the length scale of a filament. We reasoned that long-ranged and polarity independent flows may be required to maintain the proper polarity profile in spindles. Indeed, microtubule transport is necessary to counteract branching microtubule nucleation in spindles, which is known to occur in the form of autocatalytic waves that propagate away from mother microtubules (*Petry et al., 2013, Ishihara et al., 2016, Oh et al., 2016, Decker et al., 2018, Kaye et al., 2018*). Since microtubule nucleation is highest around chromosomes, in the absence of transport, autocatalytic waves naturally result in spindles with plus-ends pointing outward (*Decker et al., 2018*). Such a polarity profile is opposite to the one observed in normal spindles. A possible hypothesis is that long-range transport, enabled by gelation, is required to compensate for this inherent configuration generated by branching nucleation. Indeed, transport would be especially important near the poles where most microtubules branch off inwards and microtubule sorting is less effective (see Fig. 1F). Thus, we hypothesize that microtubule gelation might be key in maintaining the proper spindle architecture.

To test this, we simulated spindles by considering two interacting networks of opposed polarity. We included microtubule branching nucleation and we systematically varied the degree of connectivity in the network as well as the velocity of motors (Fig. 6A-D). Above the gelation point, and for sufficiently large motor velocities, we observed the emergence of a gradient of polarity similar to the one found experimentally (Fig. 6A and Movie S13), and with network densities having the same organization as in real spindles (Fig. 6D compared to Fig 1E). For additional information on the simulations, see Fig. S7 and Appendix: Branching nucleation. Strikingly, when we reduced the connectivity below the gelation point while keeping motor activity constant, filaments self-organized with a reversed polarity profile, with autocatalytic waves of filaments growing towards the poles (Fig. 6B). Similarly, when we reduced the motor activity in a gelated system, we observed the same polarity reversal (Fig. 6C).

**Figure 6:**
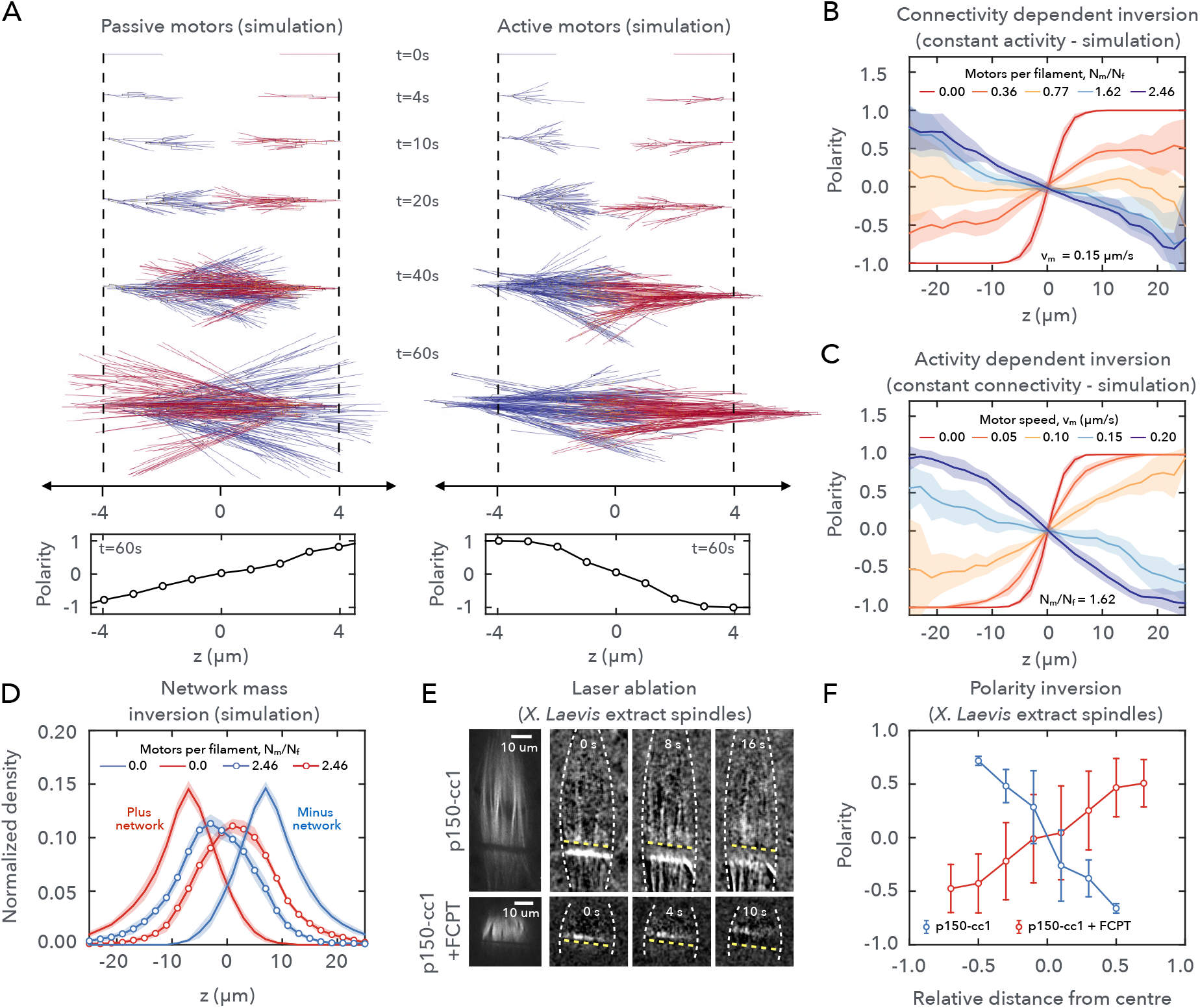
Microtubule gelation is required for the proper spindle architecture. (A) Computer simulations reveal a mechanism for generating polarity profiles observed in p150-cc1 treated *Xenopus* extract spindles. Simulation snapshots showing the time evolution of two branching filament networks, interacting via passive or active motors. In passive systems, polarity profiles are inverted compared to the control system. In active systems, local polarity sorting, and gelation lead to the establishment of control polarity profiles with transport in the opposite direction to the autocatalytic branching wave propagation (motor links in red, branching connections in blue). (B) Polarity inversion enabled by gelation transition in active systems (steady-state at *t* = 12 min and constant 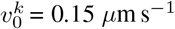). Varying connectivity by Eg5 population size (*N*_CL_/*N*_MT_ = 0.0, 0.36,0.77,1.6,2.5 from red to blue). (C) Polarity inversion enabled by activity in gelated networks (*t* =12 min and constant *N*_CL_/*N*_MT_ = 1.62). Varying activity by unloaded Eg5 velocity (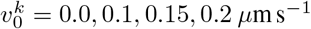 from red to blue). (D) Gelation enables the organization of control-spindle plus and minus network density organization 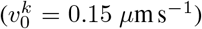. In (B), (C) and (D): branching nucleation occurs between *z* = –8 *μ*m and 8 *μ*m (see Appendix: Branching filament simulations). (E) Laser ablation for dynein inhibited spindles in the presence (see Movies S14 and S15) and absence (see Movies S16 and S17) of FCPT. Time series of differential intensity plots show depolymerization waves travelling in opposite directions, indicating inversion of filament polarity. (F) Polarity inversion in *Xenopus* extract spindles measured using laser ablation.

Our simulations predict the possibility of an ‘inverted’ spindle phenotype when motor velocity is reduced or when microtubule networks are not gelated. To test this prediction, we perturbed spindles with the Eg5 inhibitor FCPT, a drug that suppresses Eg5 ATP hydrolysis while keeping the motor strongly bound to microtubules (*Groen et al., 2008*). In spindles, this drug was shown to drastically decrease microtubule flows while maintaining a spindle-like structure (*Groen et al., 2008*). According to our simulations, spindles should undergo a microtubule polarity reversal despite resembling regular spindles. Consistently, when measuring microtubule polarity in these structures using laser ablation, we found a reversed microtubule polarity profile, with depolymerization waves propagating inwards (see Movies S14 and S15) instead of polewards (Movies S16 and S17) (Fig. 6F). Altogether these findings support that a gelation transition, together with molecular motor activity, is necessary for correct spindle self-organization.

## Discussion

Using a combination of laser ablation, speckle microscopy, and simulations, we have shown that microtubule transport in spindles can be achieved through gelation processes in two dynamic microtubule networks of opposite polarity. Our results challenge the notion that microtubule flux arises from local microtubule sorting or microtubule depolymerization at the spindle poles (*Walczak et al., 1998, Uteng et al., 2008, Loughlin et al., 2010, Needleman et al., 2010, Oriola et al., 2018*). Although the hypothesis of two independent microtubule networks was proposed in the past (*Yang et al., 2008, Takagi et al., 2019*), it remained unknown how the filaments coordinate their motion over tens of microns across the spindle. We find that microtubule gelation not only ensures force transmission and coordination along the long spindle axis but also along the short axis (see Fig. 3G), in agreement with previous results, which have shown that force transmission along the short axis helps to coordinate the poleward flows (*Shimamoto et al., 2015, Takagi et al., 2019*).

In addition to enabling long-range flows, microtubule gelation has implications on spindle mechanics. Each of the two microtubule networks is composed of aligned microtubules of the same polarity which form a crosslinked gel. We find that along their long axis these networks behave like viscoelastic fluids with a relaxation time that can be ~ 3 to 4 times larger than the microtubule lifetime. These results are in agreement with micromechanical studies of *Xenopus* spindles (*Shimamoto et al., 2011*), suggesting that long viscoelastic relaxation times might not be a consequence of the long lifetime of kinetochore microtubules (~ 5 min) but rather result from the high degree of crosslinking between microtubules. The fact that in our simulation the relaxation times of the gels do not only depend on the crosslink density but also on the mean filament length (see Fig S6B) also hints at a role of the steric interactions in long relaxation times (*Höfling et al., 2008*).

Finally, we find that a gelation process of microtubule networks, together with motor-mediated active flows, ensure the correct bipolar spindle architecture. Numerical simulations reveal that molecular motor activity, together with a high degree of dynamic crosslinking, can reverse the polarity profile that would naturally result from autocatalytic microtubule nucleation in the proximity of chromosomes. We further tested this prediction by treating *Xenopus* egg extract spindles with FCPT, a drug that suppresses Eg5 ATP hydrolysis while keeping the motor strongly bound to microtubules, and observed a reversal of microtubule polarity. Hence, our study shows the key role of gelation in spindle assembly and highlights the delicate balance between autocatalytic microtubule nucleation, microtubule transport, and crosslinking, in organizing and maintaining the proper spindle architecture. More generally, cells may use gelation of cytoskeleton structures as a physical mechanism to build and sculpt large structures made of short and dynamic components from localized motor activities.

## Material and Methods

### Cytoplasmic extract preparation, spindle assembly and biochemical perturbations

Cytostatic factor (CSF)-arrested *Xenopus* laevis egg extract was prepared as described previously (*Murray, 1991, Hannak and Heald, 2006*). In brief, unfertilized oocytes were dejellied and crushed by centrifugation. After adding protease inhibitors (LPC: Leupeptin, Pepstatin, Chymostatin) and Cytochalasin D (CyD) to a final concentration of 10 μg/ml each to fresh extract, we cycled single reactions to interphase by adding frog sperm (to 300-1000 sperm/μL final concentration) and calcium solution (10 mM CaCl_2_, 250 mM KCl, 2.5 mM MgCl_2_ to 0.4 mM Ca^++^ final concentration), with a subsequent incubation of 1.5 h. While fresh CSF extract containing LPC and CyD was kept on ice, all incubation steps were performed at 18-20 °C. The reactions were driven back into metaphase by adding 1.3 volumes of fresh CSF extract (containing LPC and CyD). Spindles formed within 30 min of incubation. We inhibited dynein with ~ 10 μM p150-CC1 (purified according to Ref. (*King et al., 2003*)) and Eg5 using ~ 100 μM FCPT. In both cases, inhibitors were added to the reactions and incubated for an additional ~ 20 min. Prior to imaging, Atto565 labeled purified porcine tubulin (purified according to Ref. (*Castoldi and Popov, 2003*)) and Hüchst 33342 were added to the reactions to a final concentration of 150 nM and ~ 16 μg/ml, respectively, to visualize microtubules and DNA. For speckle microscopy experiments, Atto565 frog tubulin was added to egg extracts to a final concentration of ~ 1 nM.

### Image acquisition

Fluorescent spindles were imaged using a Nikon spinning disk microscope (Ti Eclipse), an EMCCD camera (Andor iXon DU-888), a 60x 1.2 NA water immersion objective, and the software AndorIQ for image acquisition. The room temperature was kept at 19°C.

### Laser cutting procedure

The femtosecond laser ablation setup was composed of a mode-locked Ti:Sapphire laser (Coherent Chameleon Vision II) oscillator coupled into the back port of the Nikon spinning disk microscope and delivering 140 fs pulses at a repetition rate of 80 MHz. We used a pulse picker (APE pulseSelect) to reduce the pulse frequency to 20 kHz. Cutting was performed using a wavelength of 800 nm and typically a power of ~ 500 *μW* before the objective. The sample was mounted on a piezo stage that positioned the sample in 3D with sub-micrometer precision. The laser cutting process was automatically executed by a custom-written software that controlled the mechanical shutter in the beam path and moved the piezo stage to create the desired shape of the cut. Linear cuts were performed in several planes to cover the total depth of the spindle (~ 4-60 μm) around the focal plane. The density of cuts in depth was 2 cuts/μm to ensure proper disconnection of the spindle. Cutting was finished within 2-10 s, depending on the thickness of the cut region. Images were acquired at least every 2 s during the cutting procedure as well as for ~ 3 min after the cut. The depolymerization wave typically disappeared within 30 s.

### Simulation of active long-ranged flows

All simulations were performed using custom simulation software that is available on request. Pinning-field channel simulations are described in the Appendix: Details of the pinning field channel. Channel cross-sections are 1 *μ*m × 1 μm in (*x,y*). The domain of nucleation is between *z* = 0 *μ*m and *z* = 75 *μ*m. The average number of microtubules in the steady-state is approximately 530 with a mean length of 6.5 μm (Fig. S5). We vary the number of available motors for each state point. In the steady-state, the average number of pair bound motors will be constant. For the long, narrow channel simulations (Fig 4 D-G), the steady-state number of pair bound motors is (25, 80, 170, 315, 530, 960, 2260, 4875, 7770). For the short, wide channel simulations (Fig. 4 H-I) the range is (550, 2330, 3760). Simulation are typically run for 5 × 108 time steps with individual time steps representing Δ*t* = 5 *μ*s. We allow 1 × 108 time steps for the system to reach a steady-state before collecting data. Data is collected every 1 × 105 time steps (0.5 s). For velocity profiles, we calculate displacements every 2 s. For correlation calculations, we calculate fluctuations with time steps of 4 s. For each state point, we reduce noise by averaging over 10 simulations with the same microscopic parameters but initiated with different random seeds. We describe the details for calculating velocity profiles, connectivity plots, and correlations in detail in the Appendix. We carried out all analyses using custom MATLAB scripts.

### Active microrheology simulations

We describe the simulations to probe the material properties of our system in detail in the Appendix (Active microrheology). To avoid active flows, we set the motor velocity to zero. Otherwise, the crosslinker parameters are the same as for Eg5. For each oscillation frequency and crosslinker concentration, we reduce noise by averaging over 10 simulations. We allow 3.5 × 105 time steps before beginning the pinning field oscillations. Once the oscillations begin, simulations run for long enough that a minimum of 4 oscillation cycles can be complete. We discarded data from the first cycle to avoid transient effects. The domain of microtubule nucleation is between *z* = 0 *μ*m and *z* = 40 *μ*m. On average we have approximately 300 dynamic microtubules in the steady-state. The number of pair-bound crosslinkers is (410, 850, 1280, 1710, 2550, 5070). We set the equilibrium trap position at *z* = 20 *μ*m and the stable, trapped filament has a constant length of 7 *μ*m. We chose oscillation frequencies such that we can best probe the viscous and elastic response for the different crosslinker concentrations. Thus, we used oscillation periods of 10, 30, 45, 60, 120, 180, 300 s, all with an amplitude of *z*_0_ = 0.75 *μ*m. We recorded and analysed the trajectories for the trapped filaments using custom MATLAB scripts.

## Acknowledgments

We thank K. Ishihara for providing the construct to purify the protein p150-CC1. We acknowledge J. Pelletier and T. Mitchison for kindly providing us the FCPT inhibitor, and Martin Loose for useful discussions and feedback on the paper. We thank all the members of the Brugué’s lab for discussions and Heino Andreas for frog maintenance. We acknowledge funding from EMBO (Long-Term Fellowship with number 483-2016 to D.O.) and HFSP (CDA 74/2014 to J.B.).

## Author contributions

B.D., D.O., F.D., and J.B. conceived the project. B.D. performed computer simulations and theoretical research. D.O. performed theoretical and experimental research and analyzed the experimental data. F.D. performed experimental research and analyzed the experimental data. All authors drafted the manuscript. J.B. and F.J. directed the project.

## Supplementary Figures

**Figure S1:**
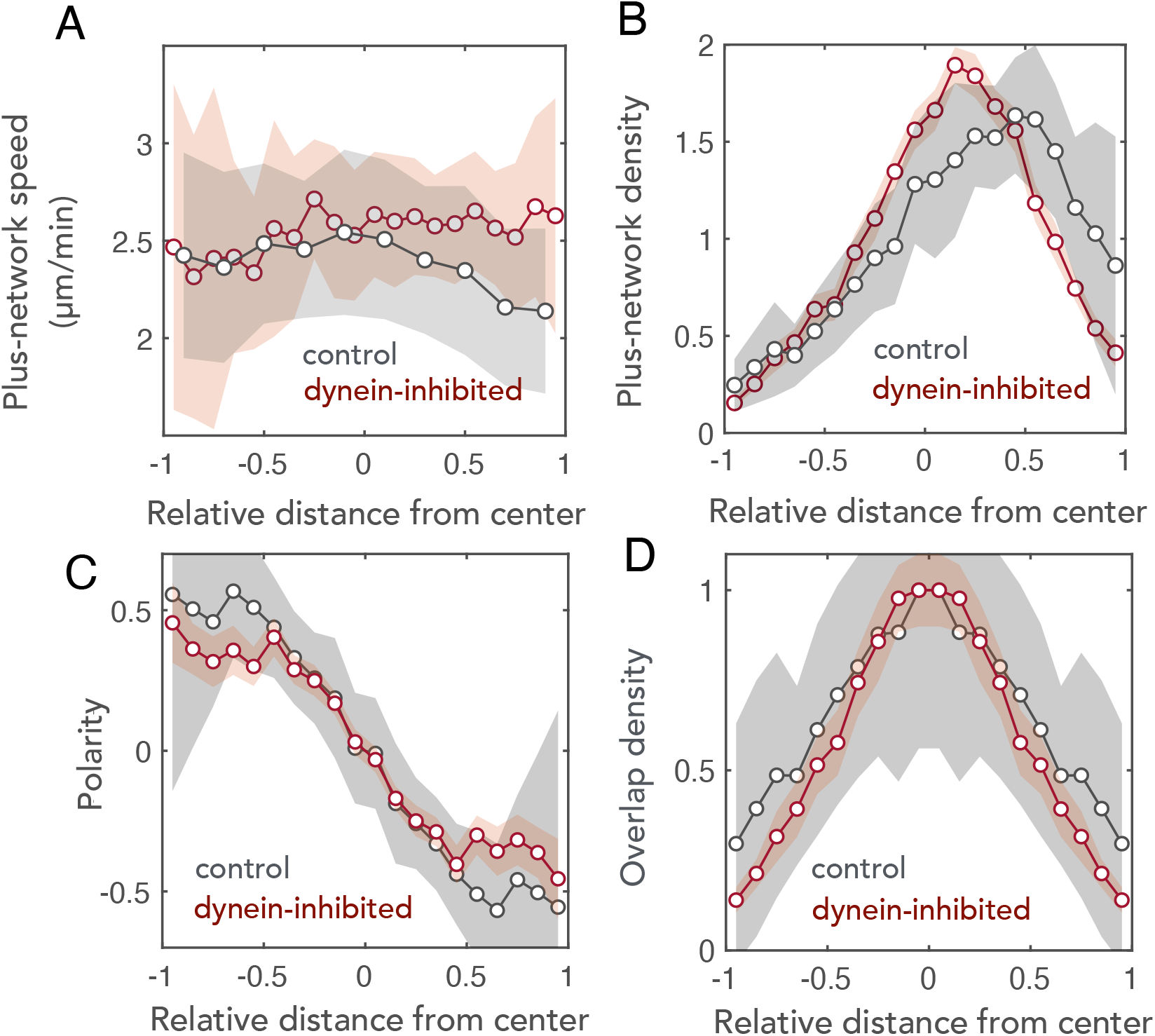
Comparison between the microtubule architecture of control (*n* = 13 spindles, black) and dynein-inhibited spindles (*n* = 10 spindles, red; data from Fig. 1). (A) Averaged velocity profiles of the plus-network (B) Density profiles of the plus-network. (C) Polarity profiles (D) Antiparallel overlap densities. Gray shaded areas indicate SD.

**Figure S2:**
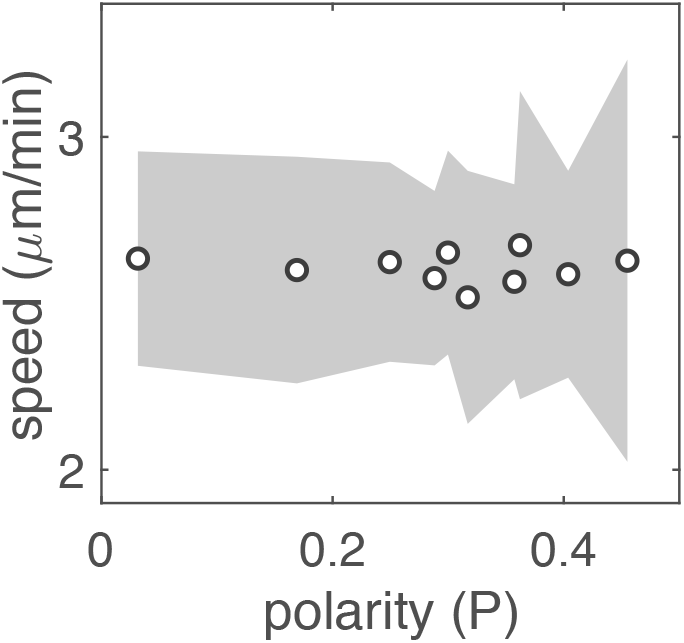
Microtubule transport velocity vs polarity using the data in Fig. 1 (F), (G). Shaded error bars correspond to the standard deviation.

**Figure S3:**
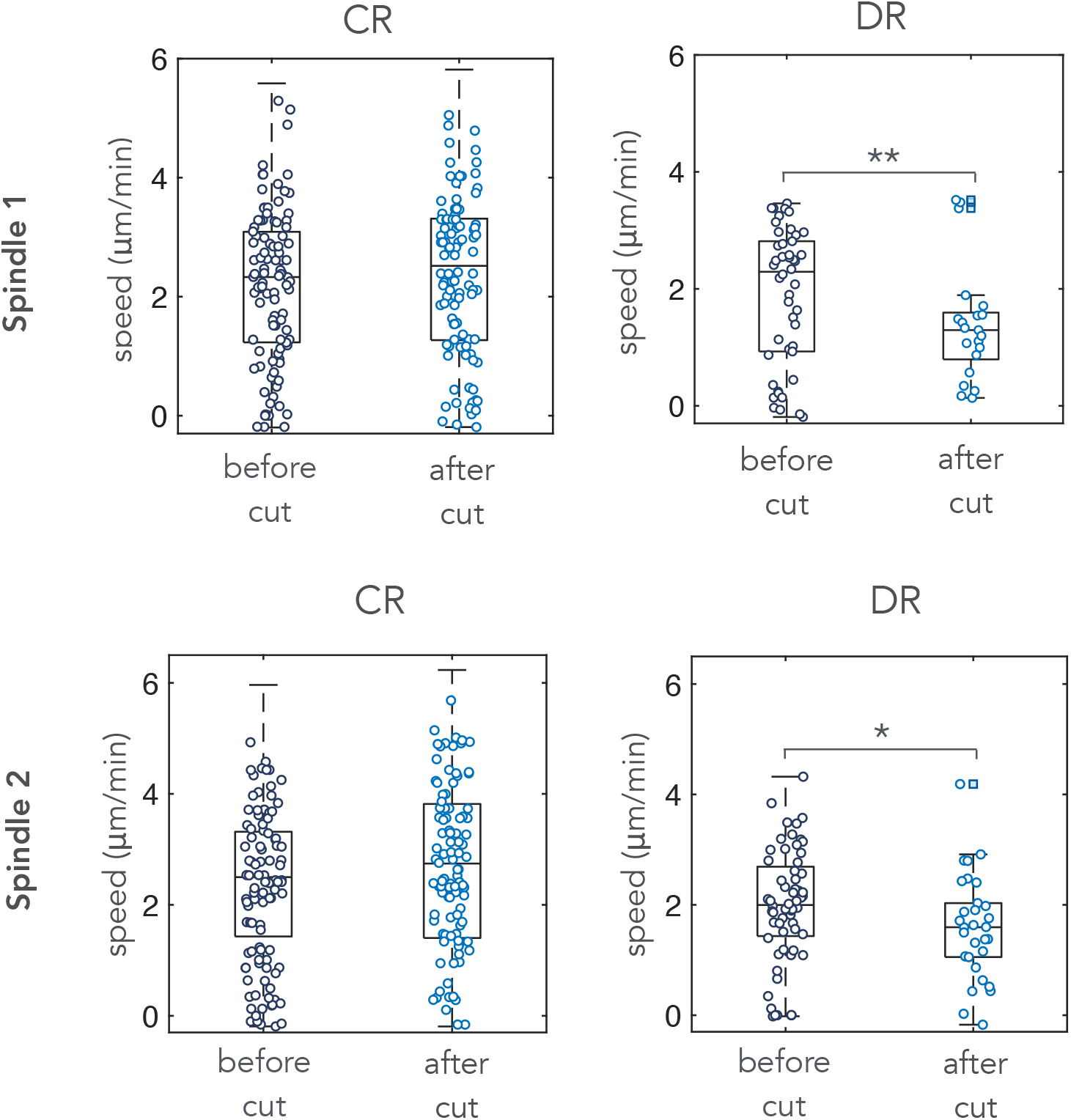
Box plots of the average on-axis velocity of speckles before and after cuts for connected (CR) and disconnected (DR) regions. Only speckles with > –0.2 *μ*m/min were considered in the analysis. A reduced subset of 100 data points is shown in the disconnected region case for the sake of clarity. * *p* < 0.05 and ** *p* < 0.01. Squares indicate outliers. The test used was a two-sample t-test. Spindle 1 corresponds to Movie S10 and Spindle 2 corresponds to Movie S11.

**Figure S4:**
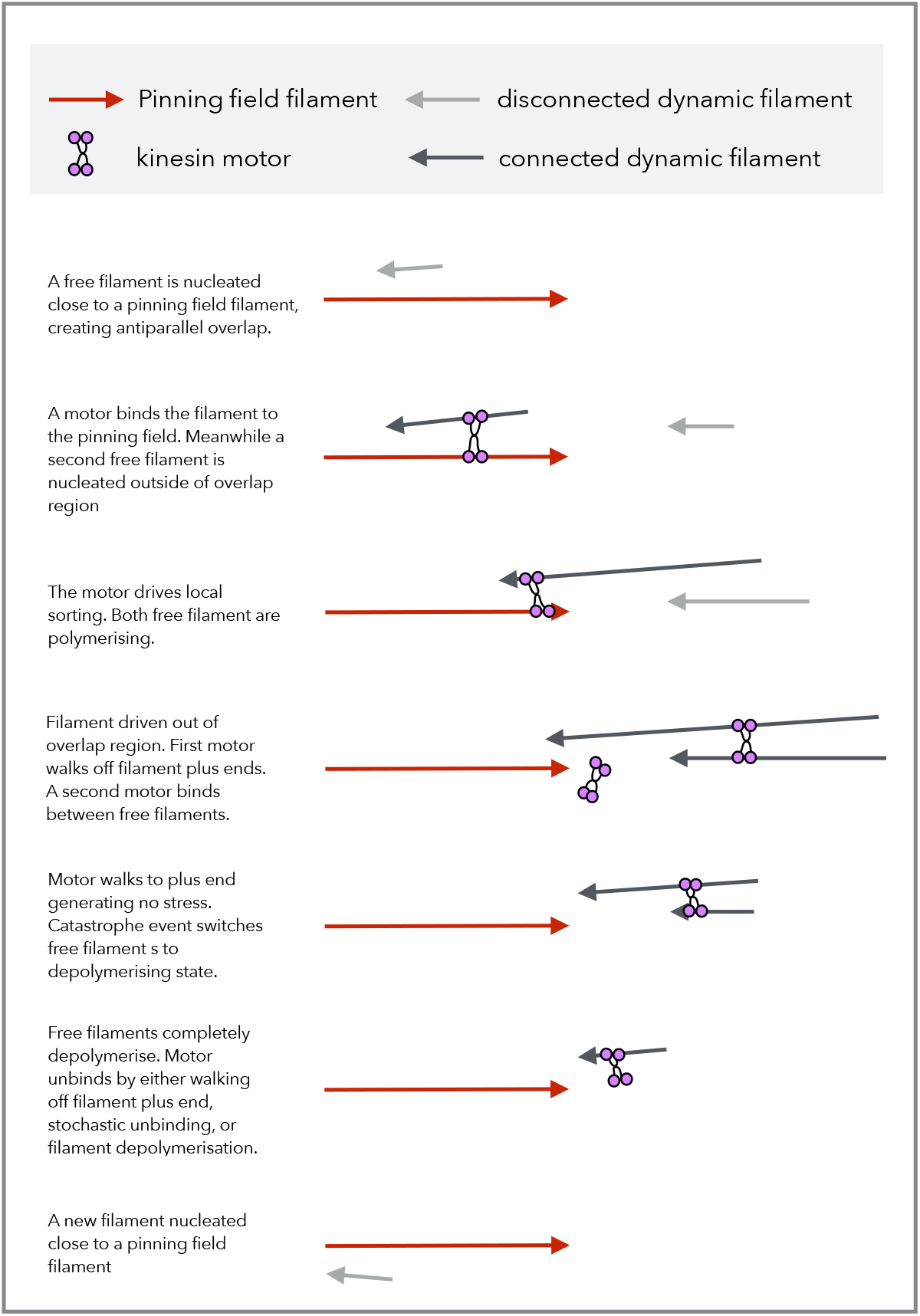
Idealize schematic showing a sequence of possible microscopic events. Emphasis is on nucleation, polymerization and depolymerization dynamics, motor dynamics and binding stochastics, as well as interactions between dynamic filaments and the pinning field.

**Figure S5:**
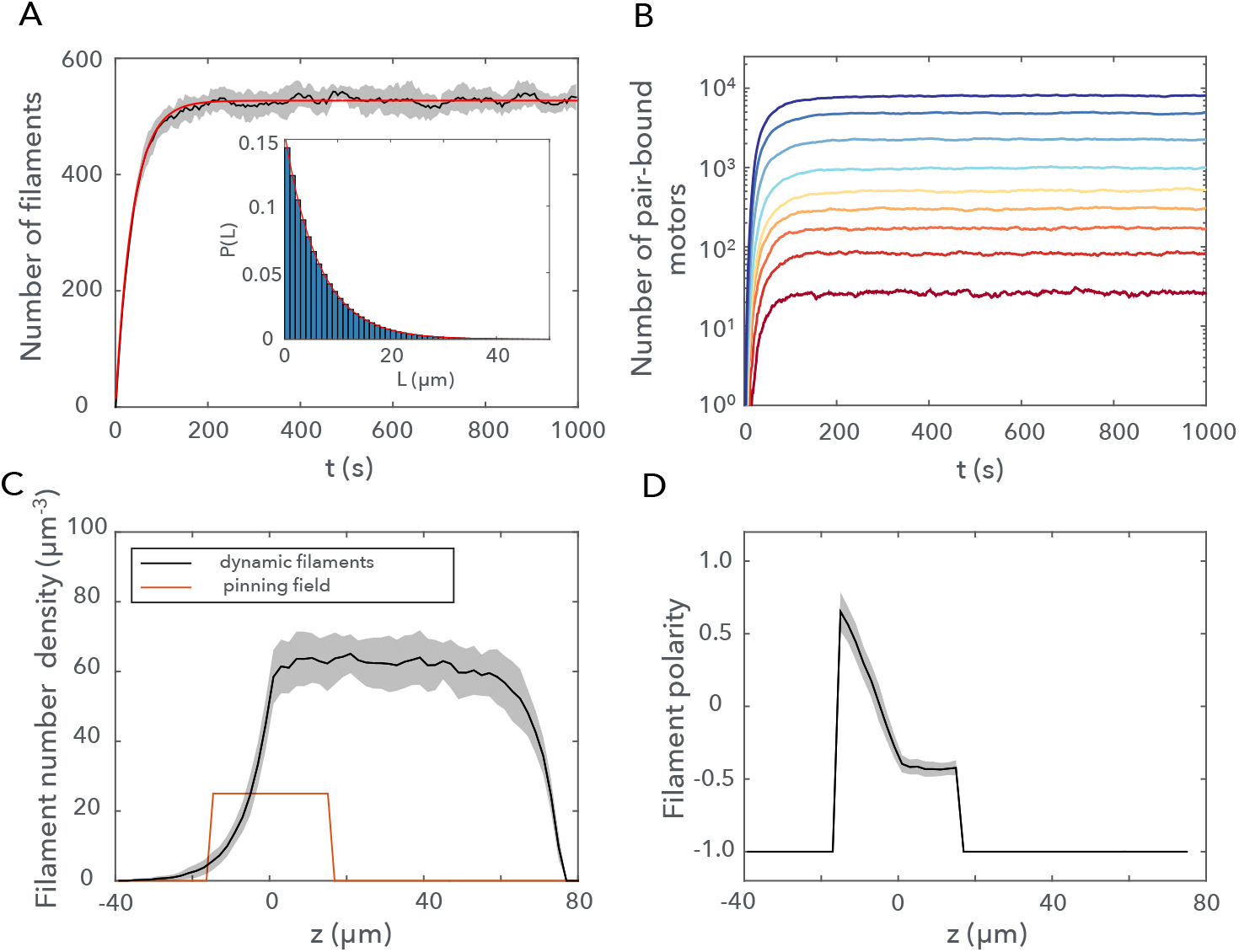
(A) Mass turnover of dynamic filaments. The trajectory in black shows 〈*N*(*t*)〉 averaged over 10 simulations with standard deviations. The red curve shows a theoretical prediction given by *N*(*t*) = *k_n_τ*(1 – exp(−*t/τ*)), where *τ* = (1 + *v_p_/v_d_*)/*k_c_* is the mean filament lifetime, *k_c_* is the catastrophe rate and *k_n_* is the nucleation rate. (inset) filament length distribution with the exponential fit (red curve) (mean length 6.5 *μ*m). (B) The number of pair-bound Eg5 motors as a function of time, corresponding to Figs. 1 (D)-(G). (C) The number density for the two distinct filament populations, calculated along the length of the channel. The finite domain of nucleation is between *z* = 0 and 75 *μ*m. (D) The total filament polarity profile, combining dynamics filaments and the pinning field.

**Figure S6:**
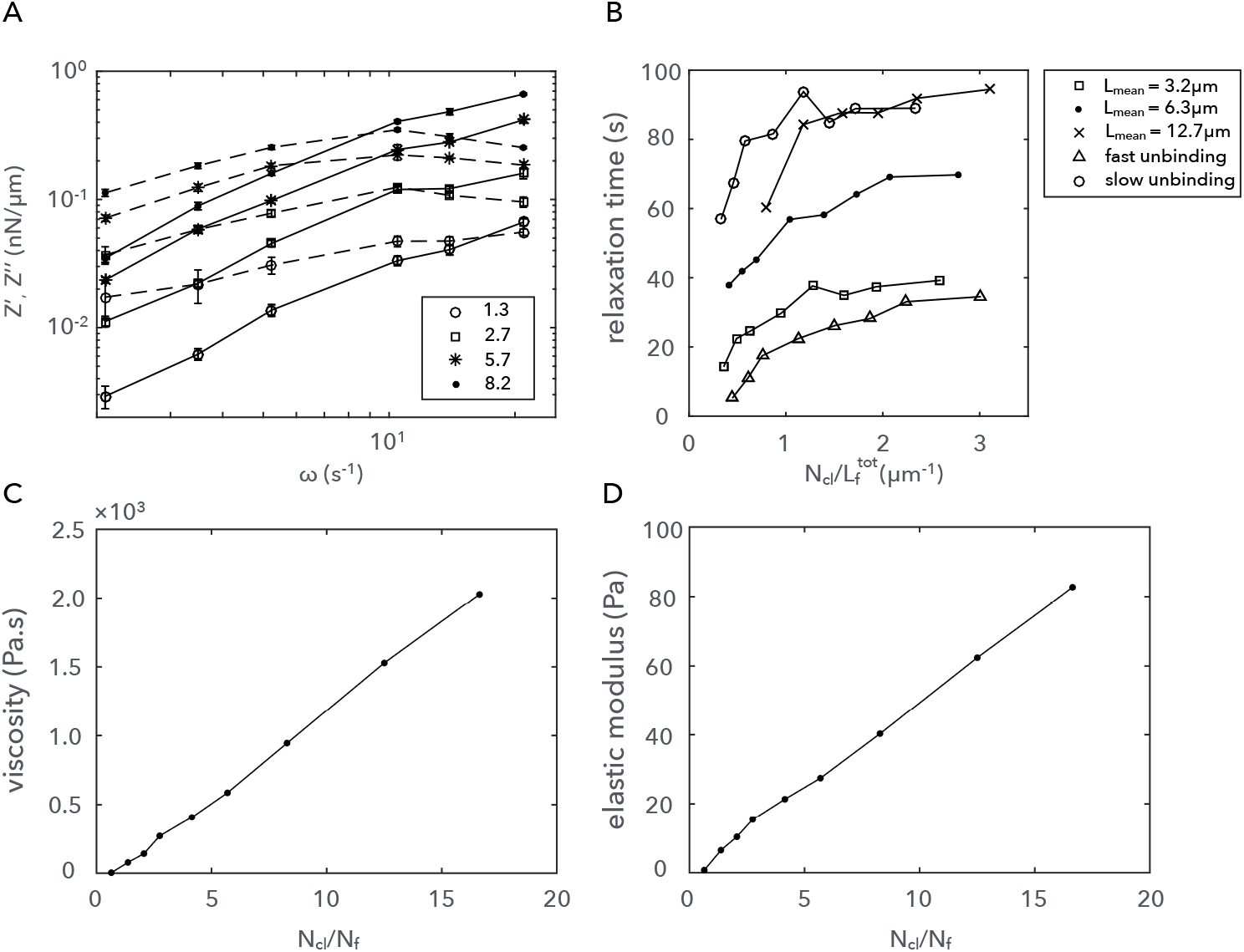
(A) Four examples of viscoelastic response, calculated with AMR. The intercept of the solid response (bold line) and the fluid response (dashed line) indicates the scaling frequency and modulus, used to scale each system for Fig. 5E. The figure legend presents *N*_CL_/*N*_MT_, as in Fig. 5E. (B) Relaxation times calculated using the AMR technique for different system parameters. The dependency on filament length distribution and the crosslinker unbinding rate is shown. The independent variable is 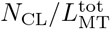 since 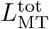 will be the same for each system with equal crosslinker concentration, whereas *N*_CL_/*N*_MT_ will vary, depending on the microtubule length distribution. Effective network viscosity (C) and elastic modulus (D) calculated using AMR (see appendix: active microrheology).

**Figure S7:**
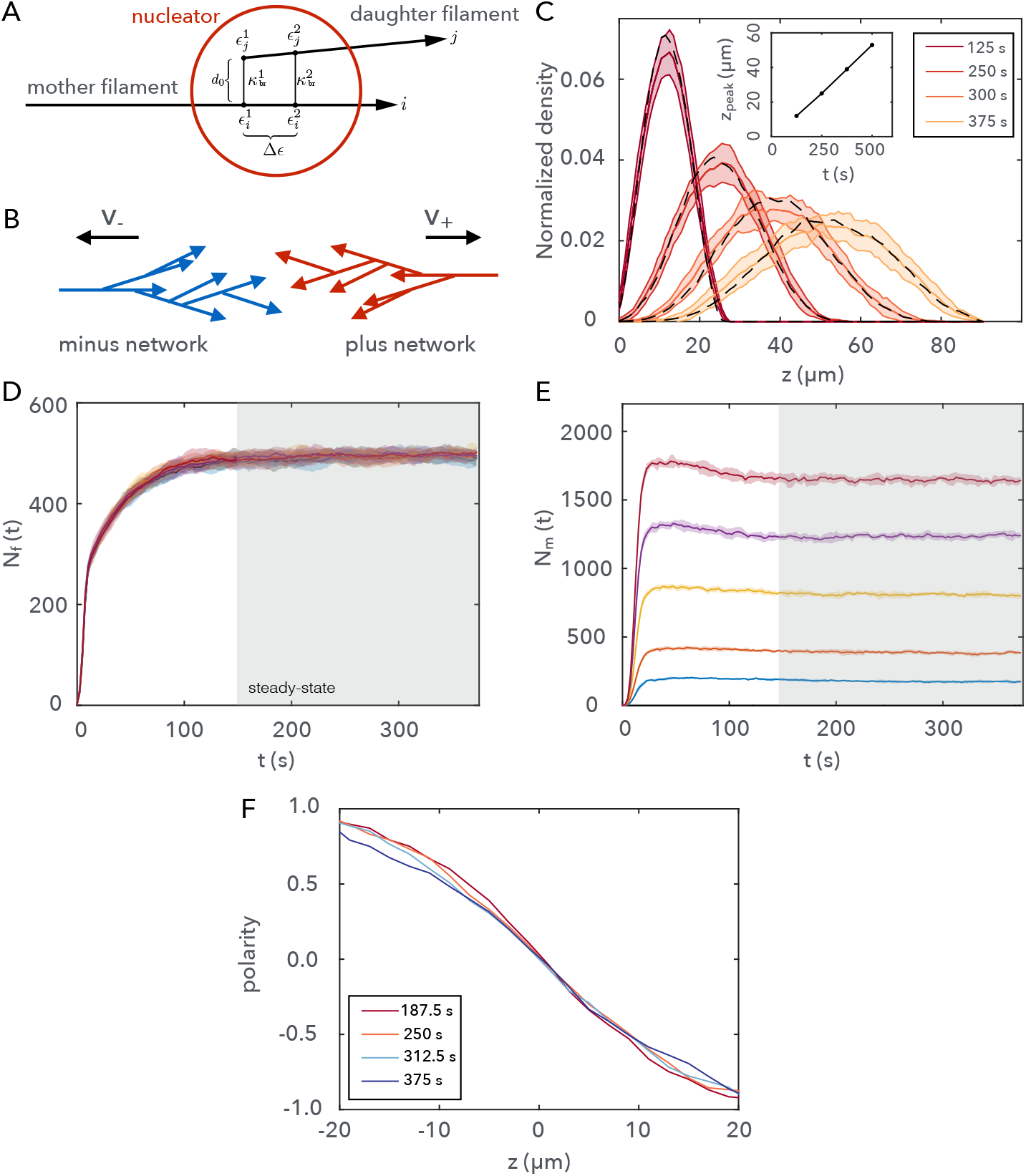
Branching filament network simulation details. (A) Schematic showing the branching connections introduced by a nucleation event. Descriptions of the various parameters are found in the Appendix: branching nucleation. (B) Plus and minus network naming conventions: naming indicates the direction of active transport due to Eg5 activity. (C) Forward propagation of a single branching network along a channel. Branching nucleation can occur in the domain 0 < *z* < 100 *μ*m. Networks begin from a single seed. A finite number of nucleators places an upper limit on the total mass density. Nucleation rate 0.1 *μ*m^−1^ s^−1^. Black dashed line for nucleation rate 1.0 *μ*m^−1^ s^−1^. (inset) mass density peak location as a function of time. The velocity of peak propagation is ~ 1.0 *μ*ms^−1^. (D), (E), and (F) details of the system steady-state for data shown in the main text Fig 6. (D) Evolution of the number of filaments for Fig. 6 (B), (C), and (D). Grey region indicates the steady-state where the total mass saturates. E) Number of crosslinkers (Eg5) in Fig. 6 (C). (F) An example showing that the polarity profile, for the two interacting branching networks, is stable over the duration of the simulation.

## Appendix

### EXPERIMENTAL METHODS

Here we provide a detailed description of the analysis of the microtubule speckle microscopy data.

#### Speckle Tracking

Microtubule speckles were tracked using the TrackMate plugin from Fiji [1]. Prior to tracking, spindles where rotated such that the long-axis corresponded to the horizontal axis of the image. We used a DoG detector with an estimated blob diameter of 4 pixels and a threshold value of ≃ 200. A simple LAP tracker was applied using a Linking and Gap-closing max distances of 4 pixels and a Gap-closing max frame gap of 2 pixels. In this way, we obtained a list of all speckles in a spindle with their corresponding spatial coordinates over time.

#### Speckle velocity

We define the position of speckle *i* at time *t* as 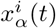 during its lifetime *t* ∈ [0, *T_i_*], where *α* = *z, y* correspond to the longitudinal and transversal components, respectively. Speckles with an absolute longitudinal velocity component smaller than 0.2 *μ*m/min were discarded from the analysis. The mean velocity components during the lifetime of speckle *i* read:

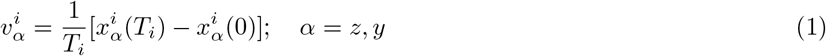

In some cases, the speckles were classified according to their direction of motion. Speckles with a longitudinal velocity larger than 0.2 *μ*m/min were included in the plus-network subset. Likewise, speckles with a longitudinal velocity smaller than if −0.2 *μ*m/min were included in the minus-network subset. The remaining speckles (≃ 4%) were discarded in the analysis. We define the position of the speckle *i* along the spindle long-axis (*z*-component) as halfway in the trajectory 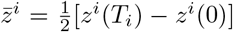. Finally, in order to compare velocity profiles between different spindles, we define the relative position of each speckle along the spindle long-axis as:

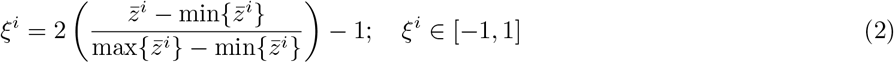

#### Speckle density profile

For each spindle *k* = 1,..., *N*, the density profiles for the plus/minus networks are formally defined as:

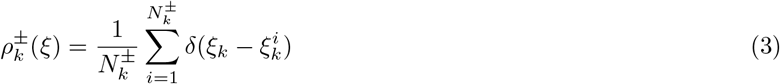

where 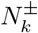 is the number of speckles in the plus/minus networks in spindle *k*. To obtain the averaged density profile we only need to compute the average for one subset (e.g. the plus-network):

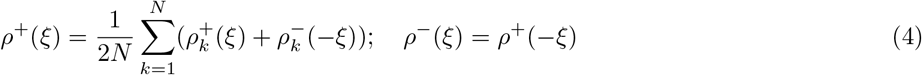

with corresponding standard deviations:

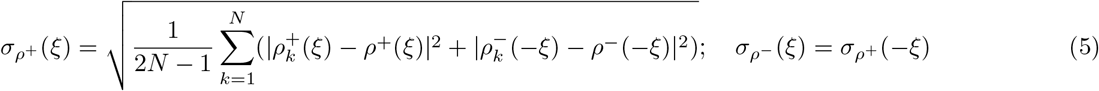

These profiles are shown in Fig. 1E in the Main Text.

#### Microtubule polarity profile

The polarity profile in Fig. 1F it is computed as:

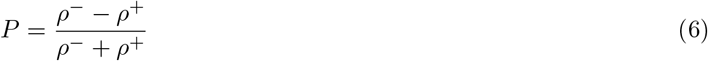

with standard deviation:

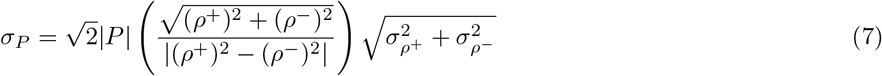

#### Microtubule overlaps profile

The profile of microtubule overlaps is computed as:

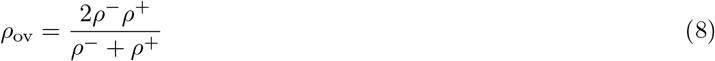

with standard deviation:

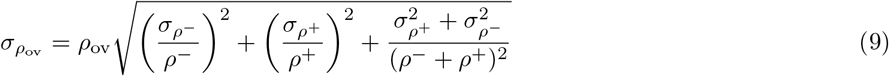

This profile is shown in Fig. 1G.

#### Microtubule velocity profile

For each spindle *k*, the velocity profiles for the networks are formally defined as:

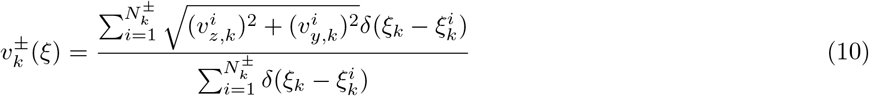

The average velocity profile in each network given *N* spindles is given by:

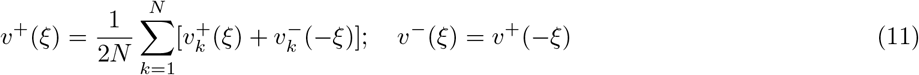

with corresponding standard deviations:

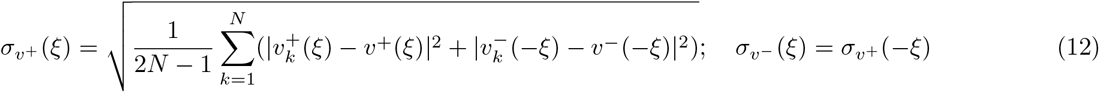

These velocity profiles correspond to the ones shown in Fig. 1-S1A. In Fig. 1G, (*v*^+^(*ξ*) + *v*^−^(*ξ*))/2 is shown.

#### Two-point microrheology

We follow an approach akin to Ref. [2] with the only difference that we substract the drift contribution to each displacement. We define the displacement of a speckle *i* along the *α*-coordinate in a timestep Δ*t* as 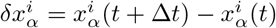. The associated relative displacement or fluctuation 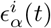 is defined as:

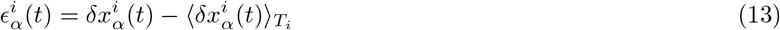

where 〈…〉 indicates a time average during the whole trajectory of speckle *i*. The equal-time two-point correlation function *C_αα_*(*r*) is defined as [2]:

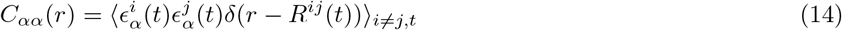

where *α* = *z,y*, 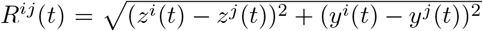 which is the distance between speckles *i* and *j* at time *t*. The average is taken over time and over all speckle pairs. Finally, the equal time two-point correlation function corresponding to shear fluctuations is computed as:

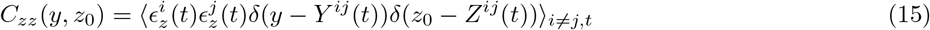

where *Y^ij^*(*t*) = |*y^i^*(*t*) – *y^j^*(*t*)| and *Z^ij^*(*t*) = |*z_i_*(*t*) – *z^j^*(*t*)|.

#### Mean Squared Relative Displacement analysis

The Mean Squared Relative Displacement (MSRD) was evaluated as follows:

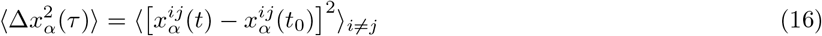

where *τ* = *t* – *t*_0_ is the time lag, *t*_0_ is the time at which the speckle pair {*ij*} first exists, 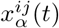 is the distance between speckles *i* and *j* in the *α*-coordinate and the average is done over all the speckle pairs.

### SIMULATION METHODS

Here we provide a detailed description of the mathematical model used for simulating cross-linked active filament networks and the method used to perform active microrheology *in silico*.

#### Filament dynamics

Filaments are represented as rigid sphero-cylinders. We define the center of mass of the *i*^th^ filament as **r**_*i*_(*t*) for *i* = 1, 2, ···, *N*^fil^(*t*), where *N*^fil^(*t*) is the number of filaments present at time *t*. **û**_*i*_(*t*) is the orientation unit vector for the *i*^th^ filament and *L_i_*(*t*) is the length of the *i*^th^ filament, which changes due to polymerization and depolymerization dynamics (see Filament mass turnover). All filaments have constant diameter *D* and anisotropic diffusion is described by three drag coefficients [3]:

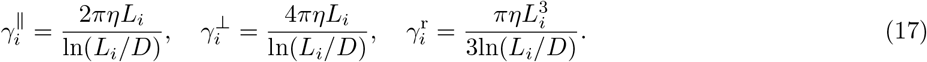

where 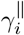 and 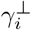 are the drag coefficients for movement parallel and perpendicular to the filament long axis, and 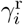 is the rotational drag for the filament center of mass. The equations of motion governing filament dynamics are [4, 5]:

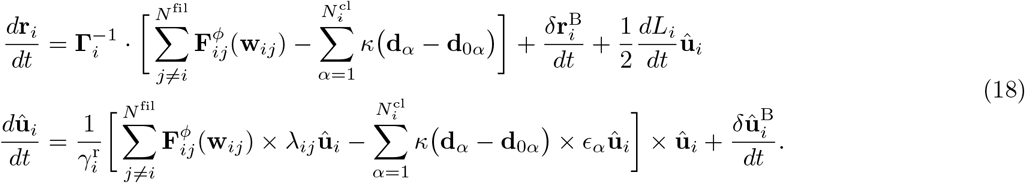

Greek indices label the connected cross-links associated with a given filament, 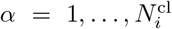. 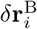 and 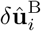 are the Brownian displacements and rotations respectively. These terms satisfy the fluctuation-dissipation theorem such that 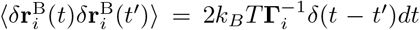 and 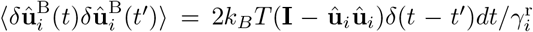, where 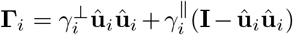 is the anisotropic friction tensor and *k_B_* and *T* are the Boltzmann’s constant and absolute temperature. 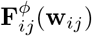 is the purely repulsive steric interaction force acting between filaments *i* and *j*. The vector **w**_*ij*_ connects two filaments and has magnitude *w_ij_* = |**w**_*ij*_|. **w**_*ij*_ is determined by treating each filament as a finite line segment and solving for a pair of line parameters (λ_*ij*_, λ_*ji*_), which locate the point on filament *i* closest to filament *j*, and *vice versa*. **d**_*α*_ is a vector connecting the two points of contact for the *α*^th^ cross-linked motor protein. The bond force is a Hookean spring with spring constant *κ* and potential *u*(*d_α_*) = –*κ*|**d**_*α*_ – **d**_0*α*_|^2^/2, where *d_α_* ≡ |**d**_*α*_| is the magnitude of the separation, and **d**_0*α*_ is the vectorial location for the equilibrium rest length along the bond direction. Finally, *ϵ_α_* is the line parameter locating the point of contact of the cross-link on the filament. The line parameter *ϵ_α_* evolves according to the rules of a given motor protein and it is subject to a two-step binding and unbinding process described below.

To calculate the magnitude of the steric interaction forces, we use a Weeks-Chandler-Anderson (WCA) force [6] with a softened core. The WCA force is a Lennard-Jones (LJ) force truncated at radial distance 2^1/6^*σ*. To soften the core, we construct a composite function with a softening parameter *B* [7]. When *w_ij_* > *Bσ*, we use the WCA force such that *σ* is the WCA radius. When *w_ij_* ≤ *Bσ* we use a linear function of *w_ij_* with a gradient given by the derivative of the WCA force at *w_ij_* = *Bσ*. The composite WCA/linear function reads:

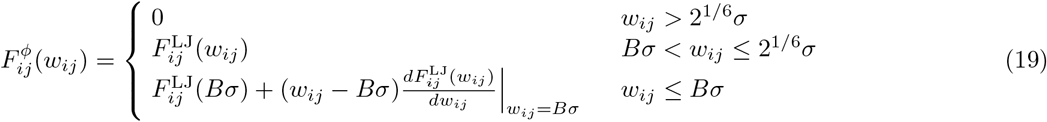

with the Lennard-Jones force given by:

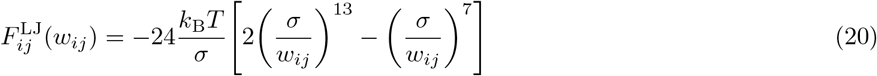

The force is then given by 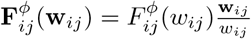.

#### Mictotubule mass turnover

We include filament dynamic instability [8] such that filaments undergo polymerization and depolymerization dynamics. We consider catastrophe events (switching from a polymerizing state to a depolymerizing state) but ignore rescue events, based on previous evidence in the spindle [9]. Polymerization and depolymerization velocities read *ν_p_* and *ν_d_*, respectively. Hence, the rate of change of the filament length in Eq. (18) follows:

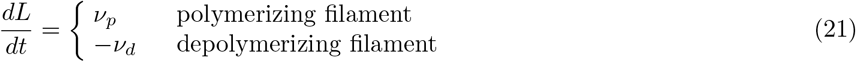

Catastrophe events occur with a rate *k_c_* and probability *k_c_*Δ*t_c_*, where Δ*t_c_* is the time interval for catastrophe sampling. For a polymerizing filament, we sample a random number *R^u^* ∈ [0,1] from a uniform distribution every Δ*t_c_* s. If *R^u^* ≤ Δ*t_c_k_c_*, then the polymerizing filament is switched to a depolymerizing state and will then depolymerize until it reaches a minimal length *L*_min_. The filament is then removed from the system.

Nucleation events occur at a rate *k_n_*. To determine the site of nucleation we draw three uniformly distributed random numbers 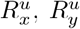 and 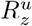 and place a new filament center of mass at 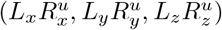 with orientation **û**_*i*_ = (0,0, –1) and filament length *L*_min_. *L_x_, L_y_, L_z_* are the simulation channel dimensions. We the bias in the orientation to ensure that we have a polar material along the channel z-axis (see Pinning field channel confinement). If the newly nucleated filament intercepts a pre-existing filament or wall (see Micro-channel confinement), we re-sample until we find a suitable position. In Fig. 4–S2A, we show the time evolution and filament length distribution of the mass-turnover algorithm.

#### Stochastic motor binding dynamics

To model motor protein binding and unbinding processes, we use a two-step model similar to Blackwell *et al*. [10]. The total number of available motors/cross-linkers is constant. Note that we typically use the symbol *N*_m_ to denote the number of active motors, and *N*_cl_ to denote the number of passive cross-linkers. In any general context, we use the term motor and cross-linker interchangeably.

Unbound motors and cross-linkers are assumed to be homogeneously distributed in a well-mixed reservoir, hence we neglect the diffusion dynamics of motors in the unbound state. For all motors in the unbound state, the probability of binding to a filament is 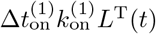, where 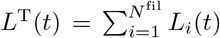 and 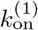 is the motor binding rate per unit length of filament. For all motors in the single bound state, the probability of unbinding and returning to the reservoir is 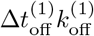. We describe a motor protein with a pair of line parameters (*ϵ_i_, ϵ_j_*) and an extension vector **d** = **d**(**r**_*i*_, **r**_*j*_, **û**_*i*_, *û_j_, ϵ_i_, ϵ_j_*). When a motor protein is in a single bound state a single line parameter *ϵ_i_* evaluates the location of the first binding domain on filament *i* and the second line parameter is inactive. The second line parameter *ϵ_j_* locates the second binding domain on filament *j* when the motor is in a pair bound state. The probability of forming (or breaking) a pair bond depends on the cross-linker extension between the two points of contact 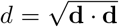. The rate of forming a pair bond to a specific location on a neighboring filament is 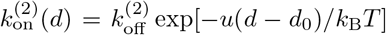, where we include the motor spring potential *u* in the Boltzmann term. The rate of unbinding depends on the force due to the cross-linker extension 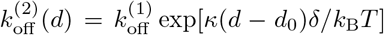, where *δ* is the scaling distance for the bond barrier crossing.

The probability that a motor will form a pair bond is calculated numerically by

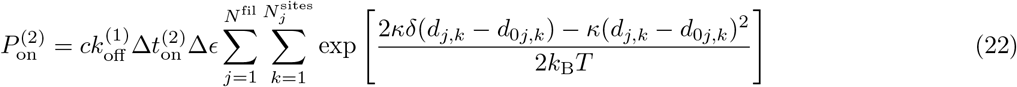

where Δ*ϵ* is the filament binding-site discretization and c represents a filament binding site concentration with units of the inverse of length. The first summation is over the discrete binding sites along the *j*^th^ neighboring filament. The second summation is over all neighboring filaments. Since the reach of the binding interaction decays rapidly, we construct a suitable neighbor list to significantly reduce the number of neighboring filaments. When a binding event is to occur, we randomly select a neighboring filament a probability weighted by each filaments contribution to Eq. 22. Subsequently, a normalized cumulative distribution for the probability density of binding (Eq. 22) along that filament is used to select the location. In Fig. 4–S2B, we show the time evolution of pair-bound cross-linkers.

#### Motor force generation

The rate at which a motor moves along a filament varies depending on whether it is under load. Motors that are in a single bound state will move at a constant speed of 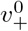. We only consider plus-end directed motors. For loaded pair-bound motors, we use a linear force-velocity relationship:

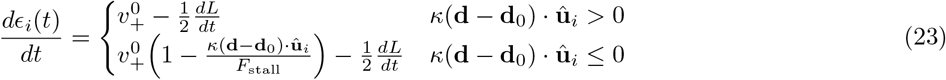

where *dL/dt* is a linear transformation due to polymerization/depolymerizatio and *F*_stall_ is the motor stall-force. In the case that *d* – *d*_0_ < 0, then 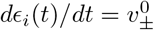.

Passive cross-linkers have zero speed and are only translated due to polymerization dynamics:

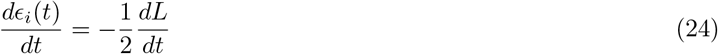

To model Kinesin-5 like motors, we define the following rules: in the event of transitioning a kinesin-5 motor from the unbound reservoir to a single-bound state, the binding location along a filament is selected from a uniform distribution biased by the total filament length. We impose a zero end-dwell-time for Kinesin-5 such that when one of the kinesin domains reaches a filament plus-end, that domain will immediately unbind. The unbinding dynamics of the second kinesin binding domain is unaffected by the unbinding of the first domain, except that there is no longer any load dependence. Likewise, if a filament depolymerizes at the location of a kinesin-5 binding domain, that domain unbinds while the other domain is unaffected.

Passive cross-links follow the same rules except that the only way they can reach a filament plus end is if the filament depolymerizes past its location.

#### Micro-channel confinement

We simulate filament networks is a micro-channel confinement geometry, with hardcore repulsive interactions between the filaments and the micro-channel walls. There are two reasons for choosing this geometry. First, the channel geometry ensures that filaments orient along a preferred axis, thus constraining the filament directionality. Second, it allows us to simulate high-density conditions with a reasonable number of filaments, thus not compromising the computational time. Given a *L_x_* × *L_y_* × *L_z_* micro-channel (with *L_z_ L_x_, L_y_*), we include hard-boundary conditions at *x* = {0, *L_x_*} and *y* = {0, *L_y_*} and open boundaries along the *z* axis. We implement filament-wall interactions in the same way as filament-filament steric interactions. For each filament, we calculate a line of closest-approach for each wall, with each wall treated as a planar surface. Four filament line parameters 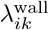 for *k* = 1, 2, 3, 4 will evaluate the points of contact on the *i*^th^ filament. We then use the distance to evaluate a composite WCA/linear function interaction 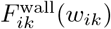, as described above. This gives four wall forces per filament

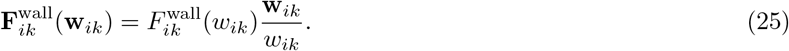

Additional force terms for the filament equations of motion (Eq. (18)) are 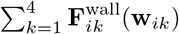, and additional torque terms are 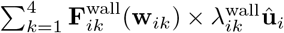. We construct and update a wall-filament neighbor list to reduce the number of potential interactions.

Additionally, we include a pinning field of stable filaments. The pinning field is an array of rigidly constrained, constant-length filaments positioned with center of mass *z*-component at *z* = 0 *μ*m. We arrange the x and y components in a periodic square lattice and we set the orientation for all filament to 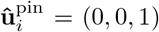. The equations of motion for the pinning field filaments are given by 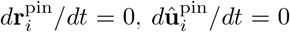 and 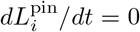. Dynamic filaments interact with the pinning field filaments via both steric interactions and motor interactions. Since dynamic filaments are always nucleated with a polarity that is opposite to the pinning field, antiparallel overlaps are restricted to the pinning field region. In Fig. 4–S1 we show an idealized representation of possible interactions between dynamic filaments, motors and pinning field filaments. In Fig. 4–S2C-D we show the density and the polarity profile for steady state mass turnover in the pinning field channel.

#### Branching nUcleation

We use a one-step model with a well-mixed reservoir of *N*^u^ unbound nucleators. The rate at which a nucleator binds to a mother filament is Δ*t*_br_*k*_br_*L*^T^(*t*), where *L*^T^(*t*) is the total instantaneous length of all filaments. A binding/nucleation event will deplete one nucleator from the reservoir. To locate the site of nucleation, we sample a random number *R^u^* from a uniform distribution and calculate *R^u^L*^T^. We organise filaments into a list, ordered by their labelling identifications. We sum the lengths of individual filaments in the list until we satisfy the condition that 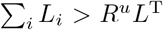. The filament label *i*, for which this inequality is first satisfied, identifies the mother filament. The location is specified by 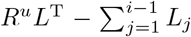, which indicates the distance from the minus-end on the *i*^th^ filament where the nucleator is placed. To bind the newly created daughter filament to the mother filament, we use a combination of two spring tethering forces located at the minus end of the daughter filament and a distance of Δ*ϵ^br^* ≈ 0.1*μm* from the minus end. By combining two tethering connections, we can restrain the rotational fluctuations of the short daughter filament. By tuning *ϵ^br^* and the spring stiffness (*κ_br_*) of the tethering connections, we can regulate the angular distribution of the resultant branching networks. We place the newly generated daughter filament at a distance of *d*_0_ from the mother filament, with a parallel configuration. We place the minus-end located tether point for the daughter filament adjacent to the selected nucleation site on the mother filament. In Fig. 6–S1A we show a schematic for the branching nucleator configuration. The tethering connections will remain fixed until either the mother or daughter filament depolymerizes past the location of the connections. To stabilize the populations of the two filament networks in the confinement channel simulations, we allow each filament population access to half of the nucleators. In Fig. 6–S1A-F we show additional results for the branching nucleation algorithm.

#### Active microrheology

We actively probe the rheological properties of cross-linked filament networks by oscillating the pinning field along the *z*-direction with period *T*. A filament probe is embedded in the network and trapped in a harmonic potential with trap stiffness *k*_trap_. To avoid actively driven flows introduced by motor activity, we use passive cross-links. The *z*-component center of mass trajectory for the pinning field is *z*_pin_(*t*) = *z*_0_sin(*ωt*), with angular frequency *ω* = 2*π*/*T*. The filament probe has a constant length and its position along the *z*-axis reads *z*_trap_(*t*). We set the equilibrium position of the trap at a distance of ~ 2-3 times that of the mean length of the dynamic filaments. We allow mass turnover and the cross-linker states to reach a steady state before we start oscillating the pinning field. Network deformations are evaluated as:

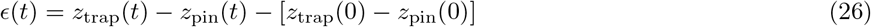

and the force response for the trapped filament is:

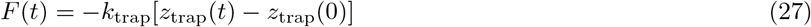

For a given frequency *ω* we calculate the in-phase (*ϵ′*) and out-of-phase (*ϵ″*) components of the deformation *ϵ*:

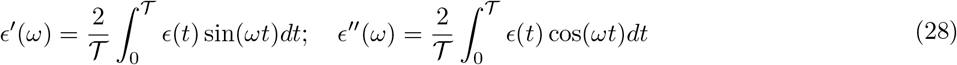

where 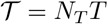 is the total time and *N_T_* is the number of oscillation cycles. Thus, we can define *ϵ*(*t*) = *ϵ*_0_sin(*ωt* + *δ_ϵ_*), where *δ_ϵ_* = arctan(*ϵ″/ϵ′*) and *ϵ*_0_ = *ϵ″*/sin(*δ_ϵ_*). Likewise, we calculate in-phase and out-of-phase components of the force response,

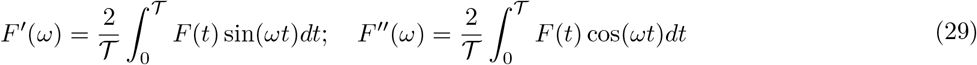

and write *F*(*t*) = *F*_0_sin(*ωt* + *δ_F_*) with *δ_F_* = arctan(*F″/F′*) and *F*_0_ = *F″*/sin(*δ_F_*). Using the relative phase shift between deformation and force response, we shift the signals and decompose such that

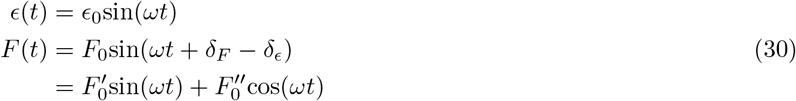

where 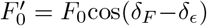 is the component of the force response signal that is oscillating in-phase with the deformation and 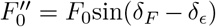 is the out-of-phase component. We define a complex response modulus *Z* = *Z′* + *iZ″*, where

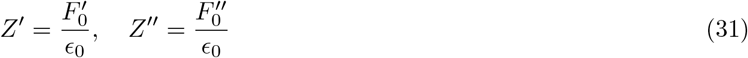

#### Calculaing network viscosity and hydrodynamic length

For a Maxwell model the frequency dependent response moduli are given by:

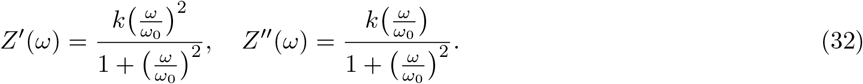

Here *k* is the network stiffness with units of Nm^−1^. *ω*_0_ = 2*π*/*τ*_0_ is the transition frequency with relaxation time *τ*_0_. From Eq. (32) we see that the network stiffness is given by *Z′*(*ω*_0_) = *Z″*(*ω*_0_) = *k*/2. To calculate the network viscosity and elastic modulus, we introduce an effective network friction coefficient:

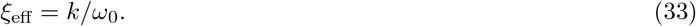

ξ_eff_ represents the drag on a filament forced through the cross-linked network. Using the effective drag coefficient for the component of motion parallel to the filament long axis Eq. (17), we evaluate the effective network viscosity:

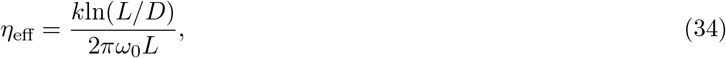

where *L* and *D* are the length and diameter of the trapped filament respectively. We can extract both *k* and *ω*_0_ from active microrheology simulations, as outlined above. The elastic modulus is then *E* = *ω*_0_*η*_eff_.

#### Velocity profiles

We track filament minus ends 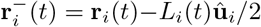 as a proxy for filament speckles. Displacements in the *z*-component 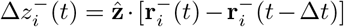 give a velocity per time increment 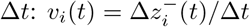. We accumulate a histogram for the filament minus-end velocities by assigning the filament velocity at time *t* to the histogram bin associated with the *z*-component displacement mid-point 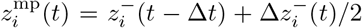. Velocity profiles are calculated in the steady-state with Δ*t* = 4 s and histogram bin width 1 *μ*m. Note that this is slightly different from the experimental approach where we use only the first and last time points of a speckle trajectory to calculate the velocity.

#### Density and polarity profiles

We construct histograms for the total filament density, the plus-network density (all filaments with *u_z_* < 0), the minus-network density (all filaments with *u_z_* > 0), and the total-polarity. The histograms represent evenly spaced bins along the *z*-axis of the simulation channel with Δ*z* = 1*μ*m. For the total-density, we identify the set of bins through which each filament passes and accumulate one count per bin. We accumulate for all filaments and then normalize by the total number of counts across all bins. For the plus and minus network densities, counting is conditioned by sgn(*u_z_*) for each filament. To calculate the polarity profiles, each filament contributes sgn(*u_z_*) to each bin through which it passes. We then divide the histogram by the pre-normalized total-density histogram.

#### Two-point microrheology

Using the minus-end trajectories 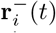, we calculate fluctuations with Eq. (13). Similarly, as in the experimental method, the correlations between minus-end speckles are calculated using Eqs.(14) and (15). The distance between two speckles is calculated in 3D as 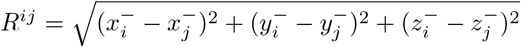, which we use to construct a histogram of fluctuation-correlations, as a function of pair separation.

#### Cluster size distribution

We calculate the distributions of cluster size using a custom MATLAB code that uses the graphconncomp MATLAB function to determine strongly connected components within networks of connected filaments. At each sample time, we create a list of all pair-bound cross-links, listed with the labels of the filaments to which it is bound. From this list, we construct a connectivity map. We pass a sparse matrix representation of this map to the graphconncomp function, which organizes a list of connected-component networks. From this, we calculate the instantaneous number of clusters and their respective sizes, which we accumulate into a histogram. The mean cluster size is 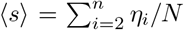 where *N* is the total number of clusters, *n* is the upper cluster size, and *η_i_* is the frequency of the *i*^th^ cluster size. The variance is normalized such that *σ*^2^ = (〈*s*^2^〉 – 〈*s*〉^2^)/〈*s*^2^〉.

### MINIMAL CONTINUUM MODEL

To understand the pinning field problem, we consider a simplified 1D continuum model. We consider a half-infinite channel immersed with a Maxwell viscoelastic fluid of viscosity *η* and relaxation time *τ*. For simplicity, we ignore bulk active stress that might be originated by dynein motors. The constitutive equation of the gel reads:

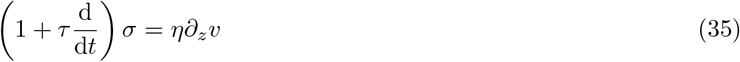

where *σ* is the stress of the gel, *v* is the velocity of the gel and 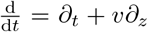. Considering the viscoelastic filament gel is permeated by a solvent of friction per unit volume *ξ*, force balance reads:

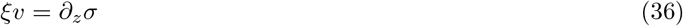

At steady state, using Eq. 36 into Eq. 35, we can rewrite Eq. 36 as:

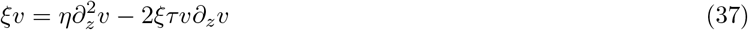

Finally the action of kinesin-5 motors is included as a stress boundary condition *σ*(0) = –*σ_e_*, where *σ_e_* > 0, while at *x* → ∞ the boundary condition is stress-free *σ*(*x* → ∞) = 0. For sufficiently small solvent friction 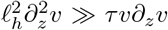, elastic effects can be neglected and we are found in the viscous fluid limit:

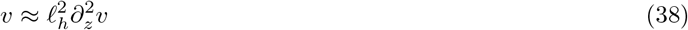

Using the boundary conditions we obtain:

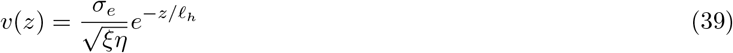

where 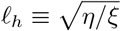 is the characteristic hydrodynamic length. We notice that for large *ℓ_h_*, the flow spans long distances. We can calculate *ℓ_h_* from simulation using the equation for filament drag in a solvent Eq. (17) and dividing by the typical volume explored by a filament, which due to the channel confinement is approximately *V* = *L_x_* × *L_y_* × *L*, for a filament of length *L*. Using *η*_eff_ measured by AMR, the hydrodynamic length in our simulation is given by

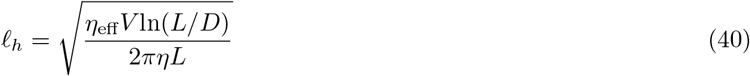

where *η* is the solvent viscosity.

### TABLE OF MICROSCOPIC PARAMETERS

**TABLE I.** A list of parameters used for simulations. References are given where appropriate. \**ϕ* = 8 nm is the size of a tubulin dimer.

| Parameter | Description | Value | Note and reference |
| --- | --- | --- | --- |
| $dt$ | simulation time-step | 2.5-5.0 μs | |
| $T$ | temperature | 300 K | |
| $\eta$ | solvent viscosity | 0.02 Pa·s | [11] |
| $D$ | filament diameter | 25 nm | |
| $c$ | filament binding sites per unit length | $1/(2\phi)$ * | order of the tubulin dimer |
| $L_{\min}$ | minimal filament length | 1 μm | $\sim 1/10$ of the typical filament length |
| $B$ | steric softening parameter | 0.9 | chosen |
| $\nu_+$ | polymerization velocity | 0.25 μm s$^{-1}$ | [9] |
| $\nu_-$ | depolymerization velocity | 0.58 μm s$^{-1}$ | [9] |
| $k_c$ | catastrophe rate | 0.038 s$^{-1}$ | set filament length distribution |
| $k_c$ | nucleation rate | 14.0 s$^{-1}$ | set filament channel density |
| $k_{\text{on}}^k$ | kinesin binding rate per unit length | $2.5 \times 10^{-4} \mu\text{m}^{-1} \text{s}^{-1}$ | Estimate |
| $k_{\text{off}}^k$ | kinesin unbinding rate | 0.03 s$^{-1}$ | [12] |
| $k_{\text{on}}^d$ | dynein binding rate per unit length | $2.5 \times 10^{-4} \mu\text{m}^{-1} \text{s}^{-1}$ | Estimate |
| $k_{\text{off}}^d$ | dynein unbinding rate | 0.1 s$^{-1}$ | [13] |
| $k_{\text{nuc}}^{\text{br}}$ | branching nucleation rate | 0.1 μm s$^{-1}$ | [14] |
| $v_0^k$ | kinesin unloaded speed | 0.05 μm s$^{-1}$ | [15] |
| $v_0^d$ | dynein unloaded speed | 0.5 μm s$^{-1}$ | [16] |
| $F_{\text{st}}^k$ | kinesin stall force | 5 pN | [17] |
| $F_{\text{st}}^d$ | dynein stall force | 1 pN | [18] |
| $\kappa_k$ | kinesin spring stiffness | 0.3 pN nm$^{-1}$ | [19] |
| $\kappa_d$ | dynein spring stiffness | 1 pN nm$^{-1}$ | [7] |
| $\kappa_{\text{br}}$ | branching tether stiffness | 0.5 pN nm$^{-1}$ | set angle distribution [14] |
| $k_B T / \delta$ | binding force scale | 3 pN | [15] |
| $k_{\text{trap}}$ | AMR trap stiffness | $5 \times 10^{-3} \text{nN } \mu\text{m}^{-1}$ | [20] |

## Notes

### Competing Interest Statement

The authors have declared no competing interest.

## References

Alvarado et al., 2013. Alvarado, J., Sheinman, M., Sharma, A., MacKintosh, F. C., and Koenderink, G. H. (2013). Molecular motors robustly drive active gels to a critically connected state. Nature Physics, 9(9):591–597.

Alvarado et al., 2017. Alvarado, J., Sheinman, M., Sharma, A., MacKintosh, F. C., and Koenderink, G. H. (2017). Force percolation of contractile active gels. Soft matter, 13(34):5624–5644.

Boukellal et al., 2004. Boukellal, H., Campás, O., Joanny, J.-F., Prost, J., and Sykes, C. (2004). Soft Listeria: actin-based propulsion of liquid drops. Physical Review E, 69(6):61906.

Broedersz and MacKintosh, 2014. Broedersz, C. P. and MacKintosh, F. C. (2014). Modeling semiflexible polymer networks. Reviews of Modern Physics, 86(3):995.

Brugués and Needleman, 2014. Brugués, J. and Needleman, D. (2014). Physical basis of spindle self-organization. Proceedings of the National Academy of Sciences, 111(52):18496–18500.

Brugués et al., 2012. Brugués, J., Nuzzo, V., Mazur, E., and Needleman, D. (2012). Nucleation and Transport Organize Microtubules in Metaphase Spindles. Cell, 149(3):554–564.

Castoldi and Popov, 2003. Castoldi, M. and Popov, A. V. (2003). Purification of brain tubulin through two cycles of polymerization–depolymerization in a high-molarity buffer. Protein expression and purification, 32(1):83–88.

Crocker et al., 2000. Crocker, J. C., Valentine, M. T., Weeks, E. R., Gisler, T., Kaplan, P. D., Yodh, A. G., and Weitz, D. A. (2000). Two-point microrheology of inhomogeneous soft materials. Physical Review Letters, 85(4):888.

Cross and McAinsh, 2014. Cross, R. a. and McAinsh, A. (2014). Prime movers: the mechanochemistry of mitotic kinesins. Nature Reviews Molecular Cell Biology, 15(4):257–271.

David et al., 2019. David, A. F., Roudot, P., Legant, W. R., Betzig, E., Danuser, G., and Gerlich, D. W. (2019). Augmin accumulation on long-lived microtubules drives amplification and kinetochore-directed growth. Journal of Cell Biology, 218(7):2150–2168.

Decker et al., 2018. Decker, F., Oriola, D., Dalton, B., and Brugues, J. (2018). Autocatalytic microtubule nucleation determines the size and mass of Xenopus laevis egg extract spindles. eLife, 7:e31149.

Dumont and Mitchison, 2009. Dumont, S. and Mitchison, T. J. (2009). Force and length in the mitotic spindle. Current Biology, 19(17):R749—R761.

Ehrlicher et al., 2015. Ehrlicher, A. J., Krishnan, R., Guo, M., Bidan, C. M., Weitz, D. A., and Pollak, M. R. (2015). Alpha-actinin binding kinetics modulate cellular dynamics and force generation. Proceedings of the National Academy of Sciences of the United States of America, 112(21):6619–6624.

Field et al., 2011. Field, C. M., Wühr, M., Anderson, G. A., Kueh, H. Y., Strickland, D., and Mitchison, T. J. (2011). Actin behavior in bulk cytoplasm is cell cycle regulated in early vertebrate embryos. Journal of Cell Science, 124(12):2086–2095.

Fürthauer et al., 2019. Fürthauer, S., Lemma, B., Foster, P. J., Ems-McClung, S. C., Yu, C.-H., Walczak, C. E., Dogic, Z., Needleman, D. J., and Shelley, M. J. (2019). Self-straining of actively crosslinked microtubule networks. Nature Physics, pages 1–6.

Gaetz and Kapoor, 2004. Gaetz, J. and Kapoor, T. M. (2004). Dynein/dynactin regulate metaphase spindle length by targeting depolymerizing activities to spindle poles. The Journal of cell biology, 166(4):465–471.

Ganem and Compton, 2006. Ganem, N. J. and Compton, D. A. (2006). Functional roles of poleward microtubule flux during mitosis. Cell cycle (Georgetown, Tex.), 5(5):481–485.

Groen et al., 2008. Groen, A. C., Needleman, D., Brangwynne, C., Gradinaru, C., Fowler, B., Mazitschek, R., and Mitchison, T. J. (2008). A novel small-molecule inhibitor reveals a possible role of kinesin-5 in anastral spindle-pole assembly. Journal of cell science, 121(14):2293–2300.

Hannak and Heald, 2006. Hannak, E. and Heald, R. (2006). Investigating mitotic spindle assembly and function in vitro using Xenopus laevis egg extracts. Nature protocols, 1(5):2305–2314.

Heisenberg and Bellaíche, 2013. Heisenberg, C.-P. and Bellaïche, Y. (2013). Forces in Tissue Morphogenesis and Patterning. Cell, 153(5):948–962.

Höfling et al., 2008. Höfling, F., Munk, T., Frey, E., and Franosch, T. (2008). Entangled dynamics of a stiff polymer. Physical Review E, 77(6):60904.

Howard, 2001. Howard, J. (2001). Mechanics of motor proteins and the cytoskeleton. Sinauer Associates.

Howard, 2009. Howard, J. (2009). Mechanical Signaling in Networks of Motor and Cytoskeletal Proteins. Annual Review of Biophysics, 38(1):217–234.

Ishihara et al., 2016. Ishihara, K., Korolev, K. S., and Mitchison, T. J. (2016). Physical basis of large microtubule aster growth. Elife, 5:e19145.

Ishihara et al., 2014. Ishihara, K., Nguyen, P. a., Groen, A. C., Field, C. M., and Mitchison, T. J. (2014). Microtubule nucleation remote from centrosomes may explain how asters span large cells. Proceedings of the National Academy of Sciences, 111(50):17715–17722.

Joanny et al., 2007. Joanny, J.-F., Jülicher, F., Kruse, K., and Prost, J. (2007). Hydrodynamic theory for multi-component active polar gels. New Journal of Physics, 9(11):422.

Kapitein et al., 2005. Kapitein, L. C., Peterman, E. J., Kwok, B. H., Kim, J. H., Kapoor, T. M., and Schmidt, C. F. (2005). The bipolar mitotic kinesin eg5 moves on both microtubules that it crosslinks. Nature, 435(7038):114–118.

Kaye et al., 2018. Kaye, B., Stiehl, O., Foster, P. J., Shelley, M. J., Needleman, D. J., and Fürthauer, S. (2018). Measuring and modeling polymer concentration profiles near spindle boundaries argues that spindle microtubules regulate their own nucleation. New Journal of Physics, 20(5):55012.

King et al., 2003. King, S. J., Brown, C. L., Maier, K. C., Quintyne, N. J., and Schroer, T. A. (2003). Analysis of the dynein-dynactin interaction in vitro and in vivo. Molecular biology of the cell, 14(12):5089–5097.

Lau and Lubensky, 2009. Lau, a. W. C. and Lubensky, T. C. (2009). Fluctuating hydrodynamics and microrheology of a dilute suspension of swimming bacteria. Physical Review E - Statistical, Nonlinear, and Soft Matter Physics, 80(1):1–15.

Lee and Pruessner, 2016. Lee, C. F. and Pruessner, G. (2016). Percolation mechanism drives actin gels to the critically connected state. Physical Review E, 93(5):52414.

Levine and Lubensky, 2001. Levine, A. J. and Lubensky, T. C. (2001). Two-point microrheology and the electrostatic analogy. Physical Review E, 65(1):11501.

Loughlin et al., 2010. Loughlin, R., Heald, R., and Nédélec, F. (2010). A computational model predicts Xenopus meiotic spindle organization. Journal of Cell Biology, 191(7):1239–1249.

Maddox et al., 2002. Maddox, P., Desai, A., Oegema, K., Mitchison, T. J., and Salmon, E. D. (2002). Poleward microtubule flux is a major component of spindle dynamics and anaphase a in mitotic Drosophila embryos. Current biology, 12(19):1670–1674.

Marchetti et al., 2013. Marchetti, M. C., Joanny, J.-F., Ramaswamy, S., Liverpool, T. B., Prost, J., Rao, M., and Simha, R. A. (2013). Hydrodynamics of soft active matter. Reviews of Modern Physics, 85(3):1143.

Mayer et al., 2010. Mayer, M., Depken, M., Bois, J. S., Jülicher, F., and Grill, S. W. (2010). Anisotropies in cortical tension reveal the physical basis of polarizing cortical flows. Nature, 467(7315):617–621.

Mayer et al., 1999. Mayer, T. U., Kapoor, T. M., Haggarty, S. J., King, R. W., Schreiber, S. L., and Mitchison, T. J. (1999). Small molecule inhibitor of mitotic spindle bipolarity identified in a phenotype-based screen. Science, 286:971–974.

Mitchison, 1989. Mitchison, T. J. (1989). Polewards microtubule flux in the mitotic spindle: evidence from photoactivation of fluorescence. The Journal of Cell Biology, 109(2):637–652.

Mitchison and Cramer, 1996. Mitchison, T. J. and Cramer, L. P. (1996). Actin-based cell motility and cell locomotion. Cell, 84(3):371–379.

Miyamoto et al., 2004. Miyamoto, D. T., Perlman, Z. E., Burbank, K. S., Groen, A. C., and Mitchison, T. J. (2004). The kinesin Eg5 drives poleward microtubule flux in Xenopus laevis egg extract spindles. J Cell Biol, 167(5):813–818.

Murray, 1991. Murray, A. W. (1991). Cell cycle extracts. Methods in cell biology, 36:581–605.

Needleman et al., 2010. Needleman, D. J., Groen, A., Ohi, R., Maresca, T., Mirny, L., and Mitchison, T. (2010). Fast microtubule dynamics in meiotic spindles measured by single molecule imaging: evidence that the spindle environment does not stabilize microtubules. Molecular biology of the cell, 21(2):323–333.

Oh et al., 2016. Oh, D., Yu, C.-H., and Needleman, D. J. (2016). Spatial organization of the Ran pathway by microtubules in mitosis. Proceedings of the National Academy of Sciences, 113(31):8729–8734.

Oriola et al., 2020. Oriola, D., Jülicher, F., and Brugués, J. (2020). Active forces shape the metaphase spindle through a mechanical instability. Proceedings of the National Academy of Sciences, 117(28):16154–16159.

Oriola et al., 2018. Oriola, D., Needleman, D. J., and Brugués, J. (2018). The Physics of the Metaphase Spindle. Annual review of biophysics, 47:655–673.

Petry et al., 2013. Petry, S., Groen, A. C., Ishihara, K., Mitchison, T. J., and Vale, R. D. (2013). Branching microtubule nucleation in xenopus egg extracts mediated by augmin and TPX2. Cell, 152(4):768–777.

Prost et al., 2015. Prost, J., Jülicher, F., and Joanny, J.-F. (2015). Active gel physics. Nature Physics, 11(2):111–117.

Redemann et al., 2017. Redemann, S., Baumgart, J., Lindow, N., Shelley, M., Nazockdast, E., Kratz, A., Prohaska, S., Brugués, J., Fürthauer, S., and Müller-Reichert, T. (2017). C. elegans chromosomes connect to centrosomes by anchoring into the spindle network. Nature communications, 8(1):1–13.

Rieckhoff et al., 2020. Rieckhoff, E. M., Berndt, F., Elsner, M., Golfier, S., Decker, F., Ishihara, K., and Brugués, J. (2020). Spindle scaling is governed by cell boundary regulation of microtubule nucleation. Current Biology, 30(24):4973–4983.

Rogers et al., 2005. Rogers, G. C., Rogers, S. L., and Sharp, D. J. (2005). Spindle microtubules in flux. Journal of Cell Science, 118(6):1105–1116.

Roostalu et al., 2018. Roostalu, J., Rickman, J., Thomas, C., Nédélec, F., and Surrey, T. (2018). Determinants of Polar versus Nematic Organization in Networks of Dynamic Microtubules and Mitotic Motors. Cell, 175(3):796–808.e14.

Sanchez et al., 2012. Sanchez, T., Chen, D. T. N., DeCamp, S. J., Heymann, M., and Dogic, Z. (2012). Spontaneous motion in hierarchically assembled active matter. Nature, 491(7424):431–4.

Shimamoto et al., 2015. Shimamoto, Y., Forth, S., and Kapoor, T. M. (2015). Measuring Pushing and Braking Forces Generated by Ensembles of Kinesin-5 Crosslinking Two Microtubules. Developmental Cell, 34(6):669–681.

Shimamoto et al., 2011. Shimamoto, Y., Maeda, Y. T., Ishiwata, S., Libchaber, A. J., and Kapoor, T. M. (2011). Insights into the micromechanical properties of the metaphase spindle. Cell, 145(7):1062–1074.

Stossel et al., 1987. Stossel, T. P., Janmey, P. A., and Zaner, K. S. (1987). The cortical cytoplasmic actin gel. In Cytomechanics, pages 131–153. Springer.

Striebel et al., 2020. Striebel, M., Graf, I. R., and Frey, E. (2020). A Mechanistic View of Collective Filament Motion in Active Nematic Networks. Biophysical Journal, 118(2):313–324.

Takagi et al., 2019. Takagi, J., Sakamoto, R., Shiratsuchi, G., Maeda, Y. T., and Shimamoto, Y. (2019). Mechanically distinct microtubule arrays determine the length and force response of the meiotic spindle. Developmental cell, 49(2):267–278.

Thawani et al., 2019. Thawani, A., Stone, H. A., Shaevitz, J. W., and Petry, S. (2019). Spatiotemporal organization of branched microtubule networks. Elife, 8:e43890.

Thorpe and Duxbury, 1999. Thorpe, M. F. and Duxbury, P. M. (1999). Rigidity theory and applications. Springer Science & Business Media.

Tolić-Nørrelykke, 2008. Tolić-Nørrelykke, I. M. (2008). Push-me-pull-you: how microtubules organize the cell interior. European Biophysics Journal, 37(7):1271–1278.

Uteng et al., 2008. Uteng, M., Hentrich, C., Miura, K., Bieling, P., and Surrey, T. (2008). Poleward transport of Eg5 by dynein–dynactin in Xenopus laevis egg extract spindles. The Journal of cell biology, 182(4):715–726.

Verma and Maresca, 2019. Verma, V. and Maresca, T. J. (2019). Direct observation of branching mt nucleation in living animal cells. Journal of Cell Biology, 218(9):2829–2840.

Walczak et al., 1998. Walczak, C. E., Vernos, I., Mitchison, T. J., Karsenti, E., and Heald, R. (1998). A model for the proposed roles of different microtubule-based motor proteins in establishing spindle bipolarity. Current biology, 8(16):903–913.

Yang et al., 2008. Yang, G., Cameron, L. A., Maddox, P. S., Salmon, E. D., and Danuser, G. (2008). Regional variation of microtubule flux reveals microtubule organization in the metaphase meiotic spindle. The Journal of cell biology, 182(4):631–639.

## References

[1] J.-Y. Tinevez, N. Perry, J. Schindelin, G. M. Hoopes, G. D. Reynolds, E. Laplantine, S. Y. Bednarek, S. L. Shorte, and K. W. Eliceiri, “Trackmate: An open and extensible platform for single-particle tracking,” Methods, vol. 115, pp. 80–90, 2017.

[2] J. C. Crocker, M. T. Valentine, E. R. Weeks, T. Gisler, P. D. Kaplan, A. G. Yodh, and D. A. Weitz, “Two-point microrheology of inhomogeneous soft materials,” Phys. Rev. Lett., vol. 85, no. 4, pp. 888–891, 2000.

[3] M. Doi and S. F. Edwards, The Theory of Polymer Dynamics. Clarendon, Oxford, 1986.

[4] Y. G. Tao, W. K. Den Otter, J. T. Padding, J. K. G. Dhont, and W. J. Briels, “Brownian dynamics simulations of the self- and collective rotational diffusion coefficients of rigid long thin rods,” J. Chem. Phys., vol. 122, no. 24, 2005.

[5] T. Gao, R. Blackwell, M. A. Glaser, M. D. Betterton, and M. J. Shelley, “Multiscale modeling and simulation of microtubule-motor-protein assemblies,” Phys. Rev. E - Stat. Nonlinear, Soft Matter Phys., vol. 92, no. 6, pp. 1–20, 2015.

[6] J. D. Weeks, D. Chandler, and H. C. Andersen, “Role of Repulsive Forces in Determining the Equilibrium Structure of Simple Liquids,” J. Chem. Phys., vol. 54, no. 12, pp. 5237–5247, 1971.

[7] P. J. Foster, W. Yan, S. Fürthauer, M. J. Shelley, and D. J. Needleman, “Connecting macroscopic dynamics with microscopic properties in active microtubule network contraction,” New J. Phys., vol. 19, no. 12, p. 125011, 2017.

[8] T. Mitchison and M. W. Kirschner, “Dynamic instability of microtubule growth,” Nature, vol. 312, pp. 237–242, 1984.

[9] J. Brugués, V. Nuzzo, E. Mazur, and D. J. Needleman, “Nucleation and transport organize microtubules in metaphase spindles,” Cell, vol. 149, no. 3, pp. 554–564, 2012.

[10] R. Blackwell, C. Edelmaier, O. Sweezy-Schindler, A. Lamson, Z. R. Gergely, E. O'Toole, A. Crapo, L. E. Hough, J. R. McIntosh, M. A. Glaser, and M. D. Betterton, “Physical determinants of bipolar mitotic spindle assembly and stability in fission yeast,” Sci. Adv., vol. 3, no. 1, p. e1601603, 2017.

[11] M. T. Valentine, Z. E. Perlman, T. J. Mitchison, and D. A. Weitz, “Mechanical properties of Xenopus egg cytoplasmic extracts,” Biophys. J., vol. 88, no. 1, pp. 680–689, 2005.

[12] M. Uteng, C. Hentrich, K. Miura, P. Bieling, and T. Surrey, “Poleward transport of Eg5 by dynein-dynactin in Xenopus laevis egg extract spindles,” J. Cell Biol., vol. 182, no. 4, pp. 715–726, 2008.

[13] R. Tan, P. Foster, D. Needleman, and R. J. McKenney, “Cooperative Accumulation of Dynein-Dynactin at Microtubule Minus-Ends Drives Microtubule Network Reorganization,” 2018.

[14] S. Petry, A. C. Groen, K. Ishihara, T. J. Mitchison, and R. D. Vale, “Branching microtubule nucleation in xenopus egg extracts mediated by augmin and TPX2,” Cell, vol. 152, no. 4, pp. 768–777, 2013.

[15] J. Howard, Mechanics of Motor Proteins and the Cytoskeleton Sunderland, vol. 55. 2001.

[16] R. J. McKenney, W. Huynh, M. E. Tanenbaum, G. Bhabha, and R. D. Vale, “Activation of cytoplasmic dynein motility by dynactin-cargo adapter complexes,” Science (80-.)., vol. 345, no. 6194, pp. 337–341, 2014.

[17] M. T. Valentine, P. M. Fordyce, T. C. Krzysiak, S. P. Gilbert, and S. M. Block, “Individual dimers of the mitotic kinesin motor Eg5 step processively and support substantial loads in vitro,” Nat. Cell Biol., vol. 8, no. 5, pp. 470–476, 2006.

[18] A. Kunwar, S. K. Tripathy, J. Xu, M. K. Mattson, P. Anand, R. Sigua, M. Vershinin, R. J. McKenney, C. C. Yu, A. Mogilner, and S. P. Gross, “Mechanical stochastic tug-of-war models cannot explain bidirectional lipid-droplet transport,” Proc. Natl. Acad. Sci., vol. 108, no. 47, pp. 18960–18965, 2011.

[19] K. Kawaguchi and S. Ishiwata, “Nucleotide-Dependent Single- to Double-Headed Binding of Kinesin,” Science, vol. 291, pp. 667–669, 2001.

[20] Y. Shimamoto, Y. T. Maeda, S. Ishiwata, A. J. Libchaber, and T. M. Kapoor, “Insights into the micromechanical properties of the metaphase spindle,” Cell, vol. 145, no. 7, pp. 1062–1074, 2011.

